# Sliding window functional connectivity inference with nonstationary autocorrelations and cross-correlations

**DOI:** 10.1101/2024.06.18.599636

**Authors:** Jing Zhang, Stefan Posse, Curtis Tatsuoka

**Affiliations:** Department of Population and Quantitative Health Science, Case Western Reserve University, OH, United States; Department of Neurosurgery, University of New Mexico, NM, United States; Department of Medicine, Division of Hematology/Oncology, University of Pittsburgh, PA, United States

**Keywords:** Dynamic functional connectivity, sliding window correlation, variance estimation, statistical inference

## Abstract

Functional connectivity (FC) is the degree of synchrony of time series between distinct, spatially separated brain regions. While traditional FC analysis assumes the temporal stationarity throughout a brain scan, there is growing recognition that connectivity can change over time and is not stationary, leading to the concept of dynamic FC (dFC). Resting-state functional magnetic resonance imaging (fMRI) can assess dFC using the sliding window method with the correlation analysis of fMRI signals. Accurate statistical inference of sliding window correlation must consider the autocorrelated nature of the time series. Currently, the dynamic consideration is mainly confined to the point estimation of sliding window correlations. Using in vivo resting-state fMRI data, we first demonstrate the non-stationarity in both the cross-correlation function (XCF) and the autocorrelation function (ACF). Then, we propose the variance estimation of the sliding window correlation considering the nonstationary of XCF and ACF. This approach provides a means to dynamically estimate confidence intervals in assessing dynamic connectivity. Using simulations, we compare the performance of the proposed method with other methods, showing the impact of dynamic ACF and XCF on connectivity inference. Accurate variance estimation can help in addressing the critical issue of false positivity and negativity.

**Highlights:**

- We study the impact of nonstationary covariance on inference in resting-state fMRI
- We propose dual sliding window estimators for correlation and its variance
- Dynamic Z scores and confidence intervals characterize fluctuations of connectivity
- A statistical manner to determine static vs. dynamic connectivity at the inter-regional level
- Improves statistical inference accuracy compared with stationary covariance model

## 1. Introduction

Functional connectivity (FC) describes the degree of synchrony of the neural activity between different brain regions, showing how activity in one region correlates with activity in another region (van den Heuvel and Hulshoff Pol, 2010). It can be measured using correlation analysis of blood-oxygen-level dependent (BOLD) signal obtained from resting-state functional Magnetic Resonance Imaging (rs-fMRI). It has been used to identify group differences in neuropsychiatric disorders, identify prognostic or diagnostic markers for individual patients, and help preoperative mapping for patients with neurological or cognitive deficits who cannot perform tasks (Fox and Greicius, 2010).

Traditionally, FC has been assumed to be stationary across time during the resting-state, which is called static FC (sFC). However, more studies support that FC is inherently dynamic with temporal changes in both magnitude and directionality, namely dynamic FC (dFC) (Hutchison et al., 2013).

sFC between two brain regions is commonly estimated using Pearson correlation, which represents the strength of connectivity. To make statistical inferences for sFC, the estimation of variance of the static correlation is crucial. However, the variance estimation of sFC is challenging because of the complicated autocorrelation structure in fMRI time series data. Ignoring the autocorrelation in the time series data can lead to an underestimation of the variance of sFC, resulting in excessive false positivity. Different statistical methods have been developed to improve the variance estimation, such as the Bartlett Correction Factor (Bartlett, 1935; Van Dijk et al., 2010). While these methods help to remedy the variance underestimation by using a deflated effective degree of freedom, a recent study (Afyouni et al., 2019) shows that existing methods fail when the correlation is not zero, and can be severely biased. The xDF method (Afyouni et al., 2019) was proposed to account not only for distinct autocorrelation in each time series, but also for instantaneous and lagged cross-correlation between time series, outperforming all existing available variance estimators under sFC.

dFC is commonly characterized by a series of correlations obtained over time from the sliding window method. Using the windowed correlation matrices across the whole brain, a classic approach is to perform clustering analysis to classify discrete FC states in the brain (Allen et al., 2014). Summary statistics for the FC states, such as the amount of time in each state and the number of transitions from one brain state to another (Choe et al., 2017), can be related to gender, age, and neurological disorder (Damaraju et al., 2014; Hutchison and Morton, 2015; Preti et al., 2017; Yaesoubi et al., 2015). Besides clustering analysis, approaches using change point detections (Cribben et al., 2012; Kim et al., 2021; Xu and Lindquist, 2015), time-varying dynamic networks (Jiang et al., 2022), and multiplication of temporal derivatives (Shine et al., 2015) have been developed to find brain FC states. These methods focus on discovering FC states for a set of brain regions simultaneously, rather than focusing on individual pair of Regions of Interest (ROIs) levels.

For the sliding window dFC correlations at the pair of ROIs level, it is often critical to determine whether the fluctuations are due to dynamic interaction patterns or inherent random noise (Kudela et al., 2017; Maleki Balajoo et al., 2020). Statistical tests based on the variance of the estimated sliding window dFC time series have been developed to test the null hypothesis of the absence of dynamicity, where the variance of the estimated sliding window dFC time series is expected to be only due to statistical uncertainties and remain relatively small. Surrogate data are used to produce the null distribution of test statistics (Hindriks et al., 2016; Sakoğlu et al., 2010). Other test statistics, such as excursion (Zalesky et al., 2014), and novel algorithms, such as kernel-reweighted logistic regression (Maleki Balajoo et al., 2020), are developed to test the null hypothesis that the estimated dFC time series is static.

Another approach to distinguishing between the spurious versus dynamic fluctuations is quantifying the uncertainty in sliding window dFC estimates using confidence intervals. A non-parametric model-free approach that combines the multivariate linear process bootstrap method and a sliding-window technique was developed to provide dFC confidence bands and assess the uncertainty in dFC estimates (Kudela et al., 2017).

However, the method lacks open-source code for easy implementation and it does not provide standardized Z scores for the dynamic correlations.

In this work, we propose a Dual Sliding Window (DSW) method for the variance estimation of sliding window correlation considering the nonstationary XCF and ACF. This approach generalizes the xDF variance estimator for sFC (Afyouni et al., 2019) and uses different sizes of sliding windows for correlation and ACF, XCF. With the variance estimator for sliding window correlations, we can not only quantify the confidence intervals for sliding window correlations, but also derive dynamic standardized Z scores, providing a means for inferential statistics beyond the sliding window correlation point estimates.

Below, we give a brief review of the xDF variance for sFC (Section 2.1), introduce the consideration of dynamic autocorrelation and cross-correlation (Section 2.2), propose our method for the variance estimator of sliding window correlations (Section 2.3), give the framework of the statistical inference for sliding window correlation (Section 2.4), generate simulated data to evaluate the proposed method (Section 2.5), and use in vivo resting-state fMRI data analysis to illustrate our method (Section 2.6). In Section 3, we present the results from the simulation for model comparison (Section 3.1), give recommendations for window size selection (Section 3.2), and present an application result to in vivo resting-state fMRI data. Section 4 provides a discussion of the results, implications and future directions. Section 5 concludes this work.

## 2. Material and methods

### 2.1 xDF variance estimation for static functional connectivity

We are interested in the relationship between two time series *y*_1_ and *y*_2_, measured over two brain ROIs. Their measurement at each time point *t* is denoted as *y*_1,*t*_ and *y*_2,*t*_, 1 ≤ *t* ≤ *T*. If we follow the assumption of sFC, the averaged correlation between *y*_1_ and *y*_2_:

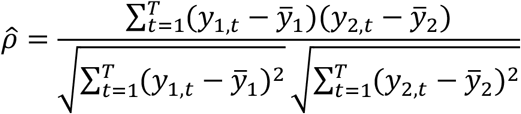

where 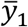 and 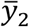 are the mean of *y*_1_ and *y*_2_, respectively.

The xDF variance estimation (Afyouni et al., 2019) for the above static 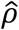 is

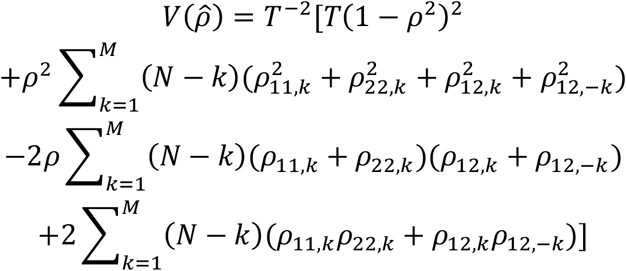

where *T* is the number of total observations, *ρ*_11,*k*_ and *ρ*_22,*k*_ are autocorrelations for two time series at lag *k*, *ρ*_12,±*k*_ are cross-correlation between two time series at positive and negative lag *k*, M is the truncation threshold for lags in autocorrelation and cross-correlation. The xDF estimator accounts for the impact of autocorrelation in each ROI, as well as the instantaneous and lagged cross-correlation. It is an improved estimator of the variance of the Pearson’s correlation coefficient (Afyouni et al., 2019), compared to the available effective degree of freedom estimators, such as Bartlett’s Formula (Bartlett, 1935), and other corrections (Bayley and Hammersley, 1946; Quenouille, 1947).

### 2.2. Autocorrelation and cross-correlation under dynamic assumption

DFC can be measured using sliding window correlations. With the window length *w*, we denote two time series starting from *t* to *t* + *w* ™ 1 as 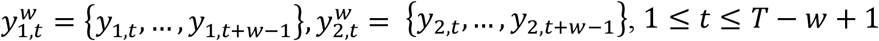 The sliding window correlation is the sample correlation coefficient between 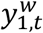 and 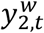 :

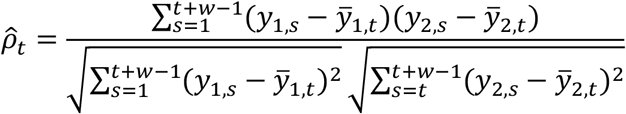

where 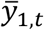 and 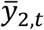 are the mean of 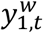 and 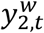, respectively.

To generalize the xDF variance estimator to the dynamic sliding window correlation, the first question is if the autocorrelation function (ACF) and the cross-correlation function (XCF) should be considered as dynamic or static. Using in vivo high-speed resting-state fMRI data (TR/TE: 400/35msec, voxel size: 3×3×3 mm^3^, number of scans: 900, for details see 2.6), we gain empirical insights into the dynamics of the ACF and XCF.

For a single subject with minimal head movement (framewise displacement < 0.1 mm), we examined the dynamic ACF at lags 1-4 of fMRI signals from Brodmann areas and the dynamic XCF at positive lags 1-4 between Brodmann areas. Figure 1a shows an example of the dynamic ACF at lags 1-4 for Bordmann area 01 in the left hemisphere (BA01L) and the dynamic XCF at positive lags 1-4 between BA01L and Bordmann area 01 in the right hemisphere (BA01R). The Pearson correlations between dynamic ACF for BA01L at lag 1 and 2, lag 2 and 3, lag 3 and 4 are 0.91, 0.94, and 0.98 respectively.

**Figure 1.**
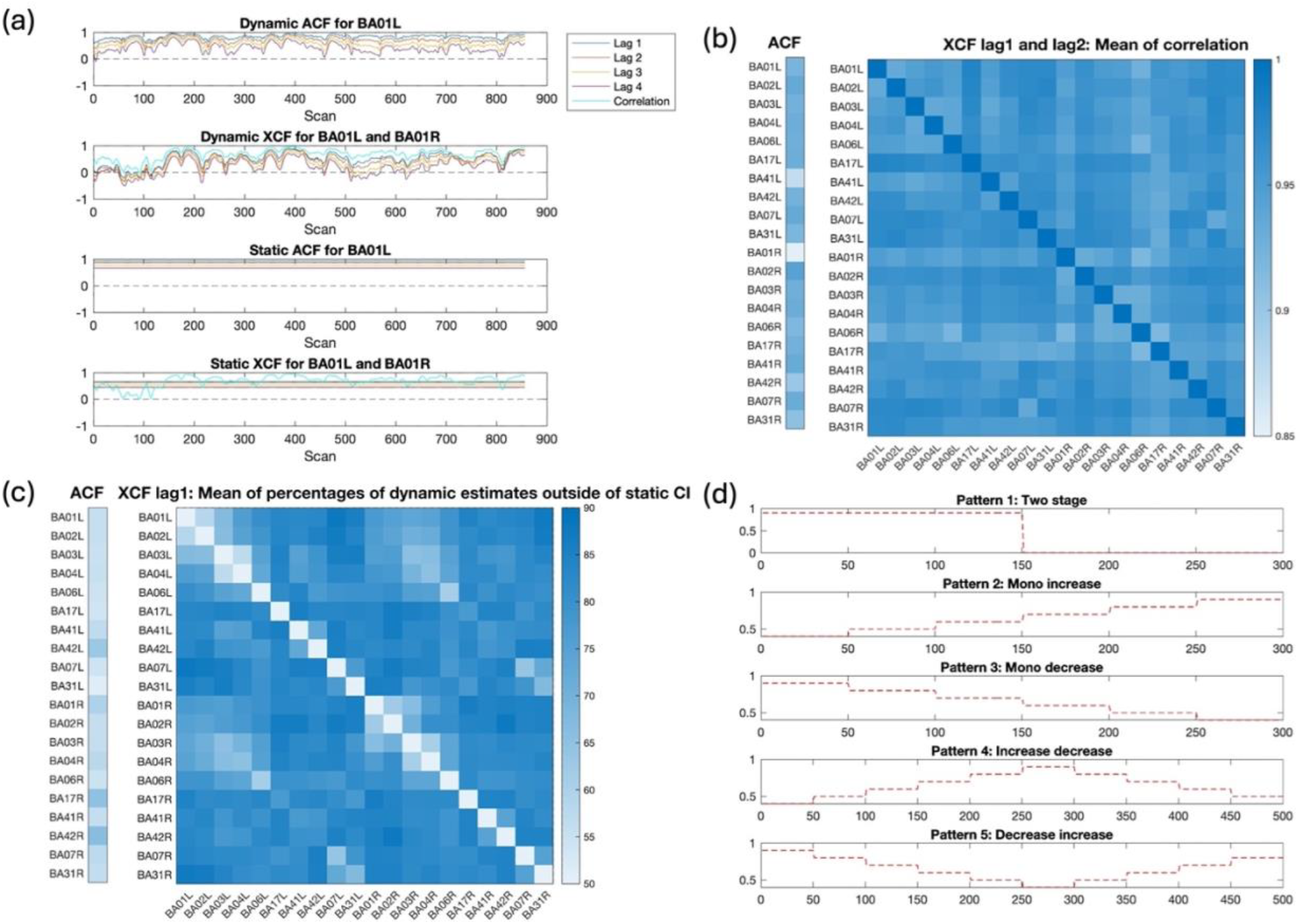
(a). Dynamic Autocorrelation Functions (ACF) and Cross-correlation Functions (XCF) compared to static ACF and XCF for a single subject’s in vivo fMRI data for Brodmann Area 1 at left (BA01L) and Brodmann Area 1 at right (BA01R). (b) Mean of correlations between ACF lag 1 and lag 2 for each ROI across subjects, and mean of correlations between XCF positive lag 1 and lag 2 for each pair of ROIs across subjects. (c) Mean of percentages of dynamic estimates outside of static CI across subjects for ACF at lag 1 for each ROI and XCF at positive lag 1 for each pair of ROIs. (d) Five dynamic correlation patterns for the simulation.

The correlations between dynamic XCF at positive lag 1 and 2, lag 2 and 3, lag 3 and 4 are 0.97, 0.98, and 0.98 respectively. These high correlations show temporal fluctuation patterns that are consistent across different lags for dynamic ACF and XCF, highlighting that the dynamic estimates are not random and there are underlying consistent dynamics in the data. In addition, the dynamic XCF estimates show a strong correlation with the dynamic correlation, which is the XCF value at lag 0, with a 0.97 correlation between XCF at lag 1 and dynamic correlations. This high correlation highlights the dynamic nature of XCF.

If we assume static ACF and XCF, we can calculate the 95% CIs of the static ACF and XCF using Bartlett’s formula (Bartlett, 1946). We then examine the percentage of dynamic ACF and XCF values that fall outside the 95% CIs of the static ACF and XCF, respectively (Supplement Figure 1). The results show that more than 50% of the dynamic estimates lie outside these intervals. This indicates the dynamic nature and significant temporal variability of the ACF and XCF.

Besides this subject’s example pair, we also look at the mean of correlations between ACF lag 1 and lag 2 for each ROI across subjects, and mean of correlations between XCF positive lag 1 and lag 2 for each pair of ROIs across subjects (Figure 1b). All means of correlations across subjects are larger than 0.85, emphasizing the consistent temporal fluctuation patterns across lags for dynamic ACF and XCF. Figure 1c shows the mean of percentages of dynamic estimates outside of static CI across subjects for ACF at lag 1 for each ROI and XCF at positive lag 1 for each pair of ROIs. All mean of percentages are larger than 50%, indicating the dynamic nature and significant temporal variability of the ACF and XCF.

### 2.3 Variance estimation of sliding window correlation

To consider the ACF and XCF within and between sliding window time series, we generalize the xDF estimator (Afyouni et al., 2019) to estimate the variance of 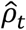 :

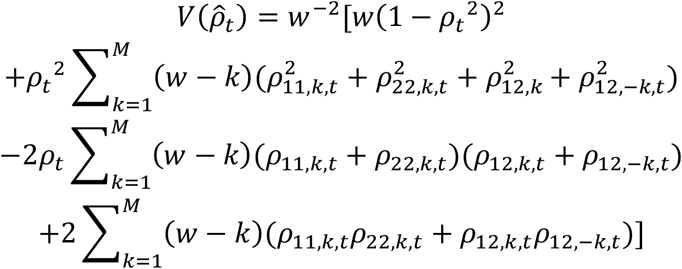

where *w* is the window length, *ρ*_11,*k*,*t*_ and *ρ*_22,*k*,*t*_ are autocorrelations for two time series at lag *k* at time *t*, *ρ*_12,±*k*,*t*_ are cross-correlation between two time series at positive and negative lag *k* at time *t*, M is the truncation threshold for lags in autocorrelation and cross-correlation.

Here, *ρ*_*t*_ can be estimated using the sliding window method with window size *w*_1_. For *ρ*_11,*k*,*t*_, *ρ*_22,*k*,*t*_ and *ρ*_12,±*k*,*t*_, we propose considering a larger window size *w*_2_ for the estimation of ACF and XCF. This is because for ACF and XCF at lag *k*, we loss *k* data points from the beginning and end of the series, reducing the effective sample size for the estimation. Hence, more data (a larger window size *w*_2_) is needed for ACF and XCF estimation compared to correlation. We refer to this method of using different window sizes during the estimation of correlation, ACF and XCF as Dual Sliding Window (DSW) variance estimator. Note that we use same window size of *w*_2_ for ACF and XCF because there is a cancellation effect between the product of ACFs and the product of XCFs at non-zero lags (Afyouni et al., 2019). Using different windows for ACF and XCF respectively appears to break this balance and result in poor variance estimation performance.

Compared with DSW, the Identical Sliding Window (ISW) variance estimator uses same window size of *w*_1_ for the estimation of ACF and XCF, so that w1 = w2. To show the importance of considering dynamic ACF and XCF for 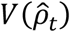, we also look at Static ACF and XCF Sliding Window (SSW) variance estimator, which assumes stationary ACF and XCF and uses the full scan of length *T* in their estimation. Note that although SSW assume stationary ACF and XCF, the correlation estimation is still dynamic.

Besides the above variance estimators based on the xDF method, we also consider the Naïve variance estimator:

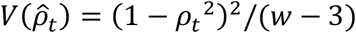

It does not consider the ACF and XCF in the data, instead assuming data independence. It is derived from the asymptotic normal distribution of the Fisher z transformation of correlation.

### 2.4 Inference for sliding window correlation

Given the asymptotical normal distribution of the Fisher z transformation of correlation, we have

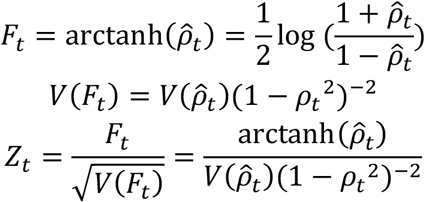

Here, *Z*_*t*_ is the standardized z score, which can be easily used as a basis for inference.

At a confidence level of 1 − α and the corresponding critical z-value *z*_1−α/2_, we can also calculate the confidence interval (CI) for *F*_*t*_ as 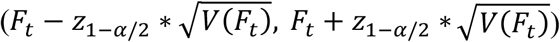 Using an inverse transformation from *F*_*t*_ to 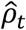

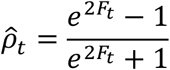

we can obtain the CI for 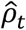 at a confidence level of 1 − α

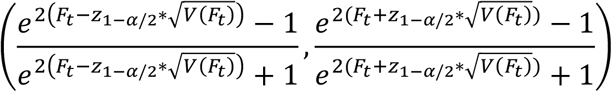

The CI width is the difference between upper CI and lower CI.

### 2.5 Simulation

In the simulation, we compare the performance of DSW against ISW, SSW, Naïve and sFC with xDF estimator, as in Section 2.1.

#### 2.5.1 Simulation setting and data generation

The objective is to simulate two time series *y*_1_ and *y*_2_ of length *T* with specific dynamic correlation, ACF, and XCF structure. We design the dynamic correlation to follow five patterns: a two stage pattern, monotonic increasing, monotonic decreasing, increase followed by decrease, decrease followed by increase (Figure 1d), similar dynamic correlation patterns considered in (Kudela et al., 2017). We use a step function for the gradual change of correlation, ensuring stationary correlation within each step. Note that length T can vary by pattern. We set the autocorrelation structure as stationary within *T* and being the same between two ROIs. Two scenarios of ACF are considered: AR4 = [0.8,0.6,0.4,0.2] and AR7 = [0.8,0.7,0.6,0.5,0.3,0.2,0.1]. The parameters are based on real data. The XCF is induced from the correlation and autocorrelation in the data.

Within a specific step of the dynamic correlation pattern, we denote the correlation for that step as *ρ*^∗^, time series within that step as 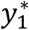 and 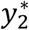 , and length of data within that step as *T*^∗^. We follow the simulation process in xDF (Afyouni et al., 2019). Assuming 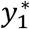 and 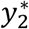 having mean zero and unit variance, the covariance matrix for 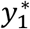 and 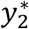 are Σ_*AC*_ and covariance matrix between 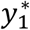 and 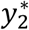 are Σ . Note that for *k* = *i* − *j*, Σ_AC_(*i, i*) = 1, Σ_*AC*_(*i, j*) = *ρ*_11,*k*_ = *ρ*_22,*k*_, representing the autocorrelation matrix, and Σ_*XC*_ (*i, i*) = *ρ*^∗^, Σ_*XC*_ (*i, j*) = *ρ*_12,*k*_, representing the cross correlation matrix.

Using Cholesky decomposition, we have 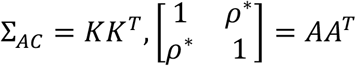, where 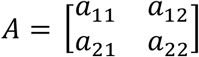. If we generate *U*∼*N*(0,1) with length 2*T*^∗^, we can set *G* = *BU*, where 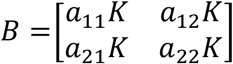. We can derive that E(*G*) = 0, 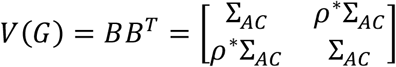. Thus, *G* is a stacked 2*T*^∗^ ∗ 1 matrix of 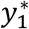 and 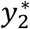, and the induced Σ = *ρ*^∗^Σ_*AC*_ . Using this framework, we can generate simulated data of 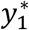 and 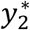 with predefined *ρ*^∗^,Σ_*AC*_ , and *T*^∗^ for each step. Connecting data from each steps during the dynamic pattern, we have the simulated *y*_1_ and *y*_2_. The simulation of each scenario is repeated 100 times.

#### 2.5.2 Evaluation criteria for methods comparison

The truth of the dynamic correlation is set in the simulation setting. However, the “truth” of the variance of sliding window correlation needs to be obtained from the simulation. For a specific time point *t*, 1 ≤ *t* ≤ *T* − *w*_1_ + 1, we obtain 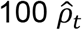 values from the 100 simulations using sliding window of size *w*_1_. The variance among these 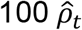 values is considered the simulated truth for the variance of sliding window correlation of size *w*_1_ at time *t*. Using true dynamic correlation and simulated true variance, we can obtain simulated true Z score and simulated true CI width as in Section 2.4.

The results are evaluated using the Mean Absolute Error (MAE) between the corresponding estimates and the truth. We choose MAE over Mean Squared Error (MSE) because MAE is less sensitive to outliers. MAE is calculated for dynamic correlation, variance, Z score and CI width at each time point. The mean MAE along the time series is calculated as an overall statistics. Smaller MAE indicates better performance. CI coverage percentages of truth correlation and the zero correlation are also calculated for each time point, representing percentage of times the true correlation or zero correlation is within the CI among 100 repetitions. Mean CI coverage percentages along the time series are also calculated.

#### 2.5.3 Window size selection for w_2_

For the DSW method, we need to select window size *w*_1_ for correlation and *w*_2_ for ACF and XCF. Selecting *w*_1_ can be based on existing discussions (Hindriks et al., 2016; Leonardi and Van De Ville, 2015; Savva et al., 2019; Shakil et al., 2016; Zhuang et al., 2020). Main factors influencing the sizes of *w*_1_ selection include temporal dynamics of resting-state networks and autocorrelation lag order. Below, we suggest a heuristic approach for selecting *w*_2_ based on *w*_1_ if one follows the window selection approach we employed. From our simulations, it appears that *w*_1_ and *w*_2_ are inherently related in that the ratio of their values falls within a consistent range. The results generated here can potentially serve as a reference and guide for future use of our method.

For each simulation scenario, we first select the size for *w*_2_ by calculating the mean MAE for XCF at positive lag 1 across various *w*_2_ window size values. Note that we use XCF at positive lag 1 as a proxy for overall XCF, since lag 1 is most important among lags. There exists a window size, 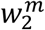 that yields the minimum mean MAE within the range of window sizes considered. From the simulations considered, the mean MAE as a function of window size is convex, which would ensure a unique minimum.

Second, we establish a tolerance percentage *q*, which permits an allowable *q*-percentage increase in MAE from the minimum mean MAE. This tolerance percentage *q* helps identify a window size that is nearly equivalent to 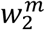 but smaller, thereby preventing an unnecessary increase in window size for a relatively small decrease in MAE. Larger window sizes with similar MAE values tend to smooth out the dynamic properties and fail to capture them effectively (Iraji et al., 2021) (see discussions for details). The MAE criterion, while insightful as it minimizes a measure of bias, appears to have a tendency to “play the middle” in terms of smoothed estimates of dynamic correlation over the time course. We thus opt for a trade-off of somewhat smaller window size at a pre-determined sacrifice of loss in MAE. We denote 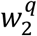 as the smallest window size that satisifies the tolerance percentage *q*. For a same simulation scenario, we similarly determine the sizes for *w*_1_, denoted as 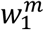 and 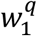 .

We summarize the relationship between 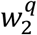 for the XCF at positive lag 1 and 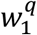 for the correlation under various simulation scenarios, same to the settings described in Section *2*.*5*.*1*. Additionally, we consider a range of *q* percentages (5%, 10%, 15%) in the simulations and examine how this relationship changes across different magnitude of *q* (small, medium, large).

### 2.6 In vivo resting-state fMRI data analysis

We use a dataset collected on a 3 Tesla clinical scanner using Simultaneous Multi-Slice Echo-Planar Imaging (SMS-EPI) in six healthy controls (Vakamudi et al., 2020). The TR/TE is 400/35msec, the voxel size is 3×3×3 mm^3^ and the number of scans is 900. The data was preprocessed using TurboFIRE software (Vakamudi et al., 2020), to perform motion correction, spatial smoothing (5mm isotropy), temporal moving average low pass filtering (4 seconds), functional anatomical segmentation in subject space into 144 Brodmann area (BA) seeds using the Talairach Daemon database, and extraction of ROI averaged time course data. Further processing in MATLAB regressed out the confounding signals including six movement parameters, averaged Cerebrospinal fluid (CSF), averaged White Matter (WM) and sine-cosine detrending for high-pass filtering up to fifth order. The following ROIs on the left and right sides were included in the analysis: BA01-03 in the sensorimotor network, BA04 and BA06 in the motor network, BA17 in the visual network, BA41 and BA42 in the auditory network, and BA07 and BA31 in the default mode network. Note that subject 5, with minimal head movement, (Supplement Figure 2) is used to examine the dynamics in ACF and XCF in Section 2.2 and 3.1.

**Figure 2.**
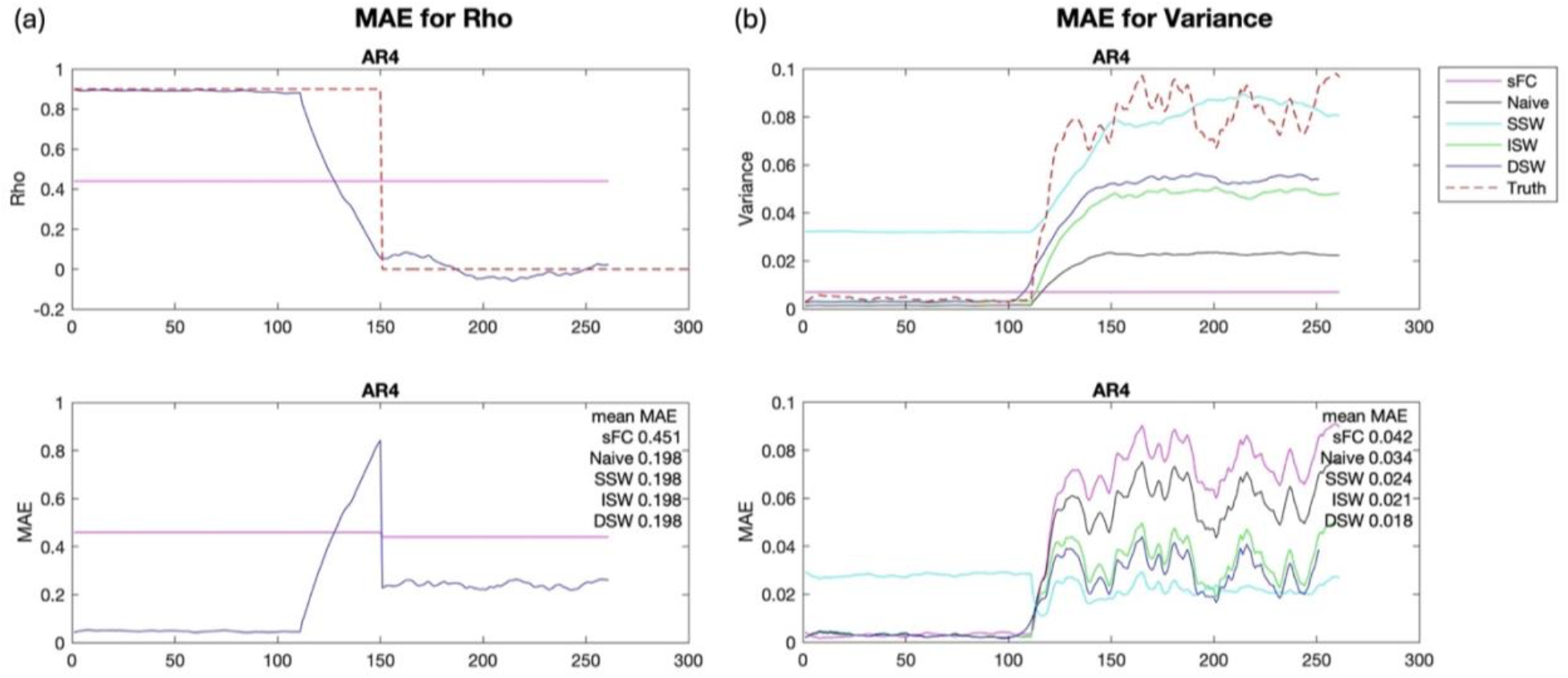
Simulation results comparing different variance estimators for simulation scenario with dynamic correlation Pattern 1 (two stage) with AR4 structure. Upper panels in each figure are the mean of 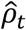 and 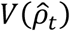 , respectively, among 100 simulations for different methods. Lower panels in each figure are Mean Absolue Error (MAE) of 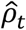 and 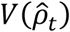 , respectively. Autocorrelation structure is autoregressive order of 4 (AR4).

For each pair of ROIs, we estimate sliding window correlation with window size (*w*_1_) 45 scans (18 seconds) and the variance of sliding window correlations using DSW method with window size (*w*_2_) 55 scans (22 seconds). In addition, we look at SSW results to compare with DSW. We also look at DSW method with *w*_1_ = 75 scans (30 seconds) and *w*_2_ = 90 scans (36 seconds) for sensitivity consideration. For each subject and pair of ROIs, we estimate the mean and standard deviation of dFC Z scores across the scan, 95% CI coverage of zero correlation, and CI coverage of sFC. Wilcoxon signed rank tests are used to compare between methods.

### 2.7 Data and code availability

The data is confidential, but will be shared upon reasonable request. The code will be available on Github.

### 2.8 Ethics statement

The SMS-EPI study received ethical clearance from the Institutional Review Board at the University of New Mexico. Written informed consent was obtained from adult participants.

## 3. Results

### 3.1 Simulation results of method comparison

In this section, we use *w*_1_ = 40, *w*_2_ = 50 when comparing between methods. The details of window size selection will be discussed in Section 3.2.

Figure 2-4 shows the simulation results for Pattern 1 two levels of correlation with AR4 covariance structure. The upper panels in Figure 2 and Figure 4 show the mean of 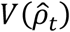 , *Z*_*t*_ and CI width, respectively, among 100 simulations for different methods. The lower panels in Figure 2 and Figure 4 show the results of MAE for 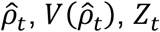 and CI width, respectively. Importantly, Figure 3 shows the percentages of dFC CI covering true correlation and zero correlation. The results are presentedalong the sliding window series in the *x* axis.

**Figure 3.**
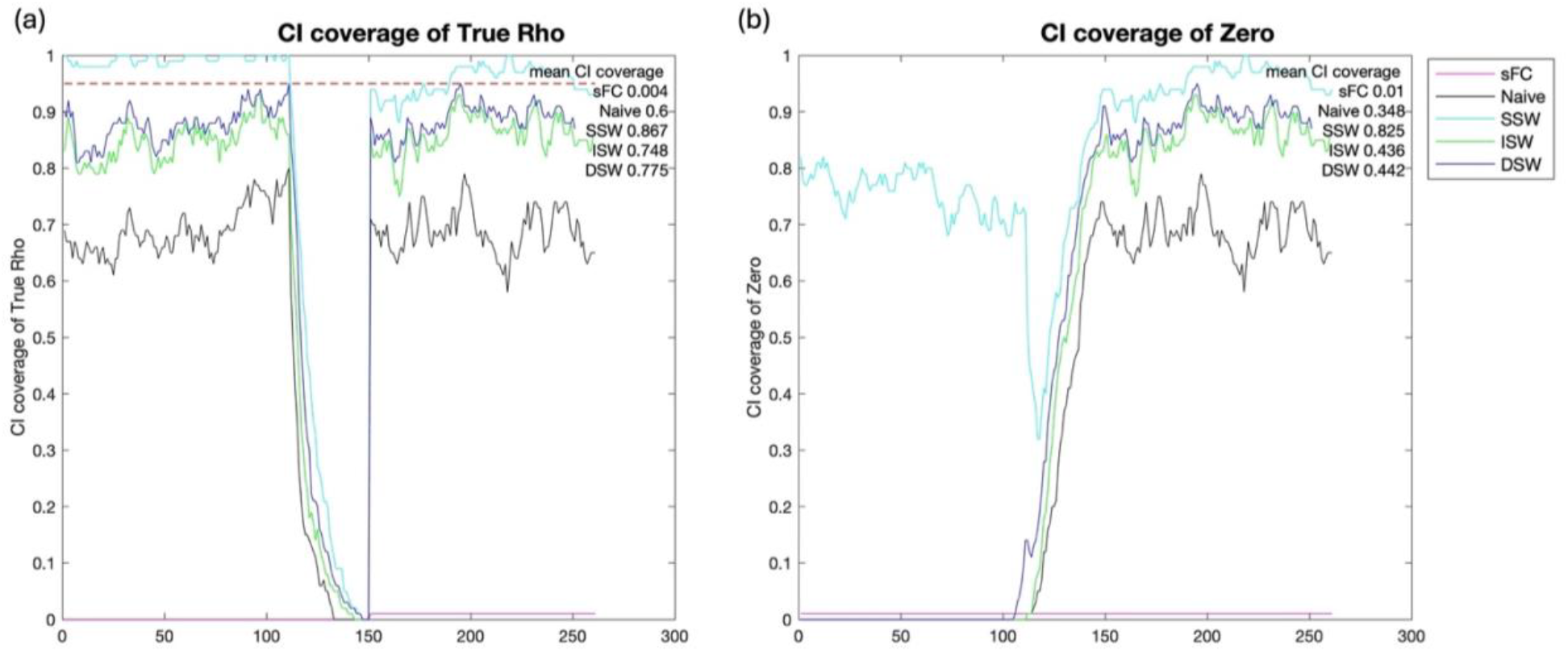
Simulation results comparing different variance estimators for simulation scenario with dynamic correlation Pattern 1 (two stage) with AR4 structure. CI: Confidence Interval. CI coverages of true rho and zero are the percentages of dynamic correlation CI covering true rho and zero correlation across simulation repetitions, respectively.

**Figure 4.**
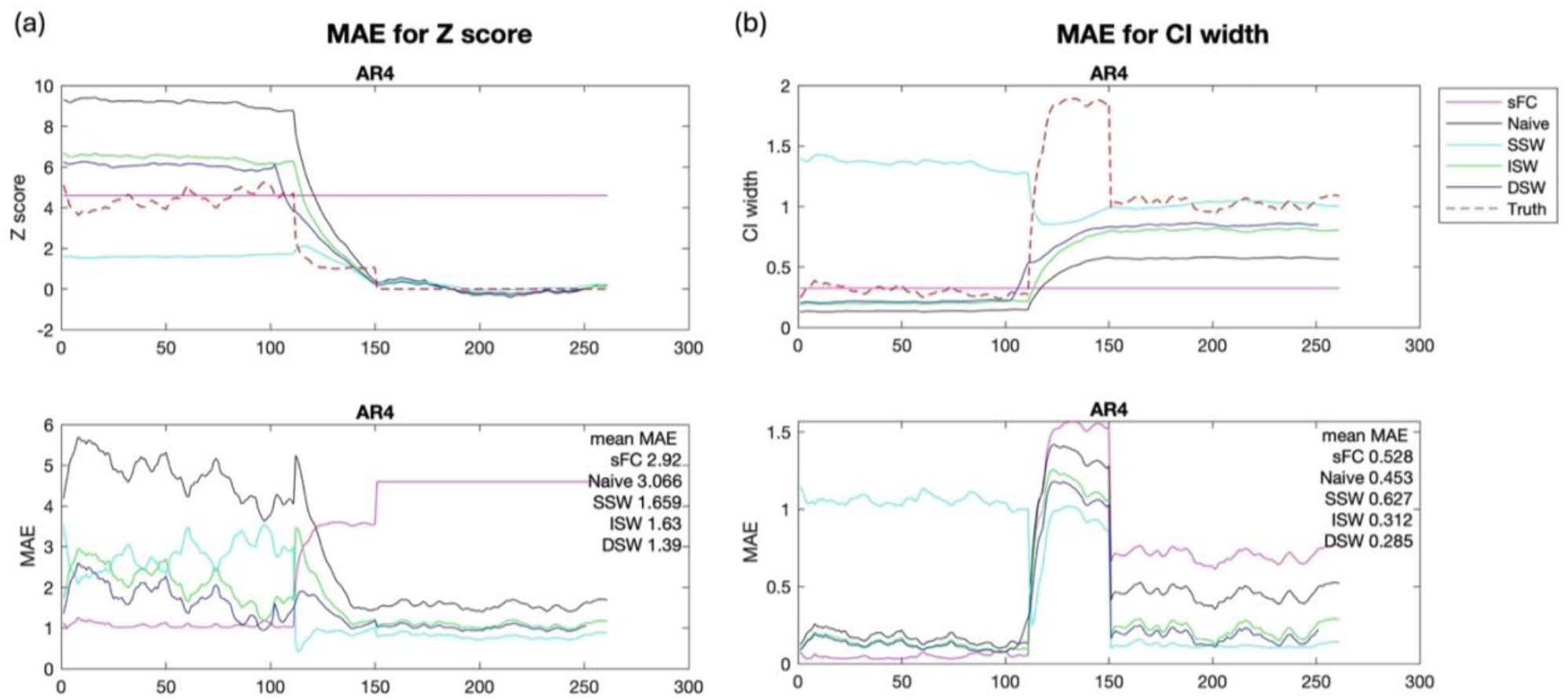
Simulation results comparing different variance estimators for simulation scenario with dynamic correlation Pattern 1 (two stage) with AR4 structure. Upper panels in each figure are the mean of *Z*_*t*_ and Confidence Interval (CI) width, respectively, among 100 simulations for different methods. Lower panels in each figure are Mean Absolue Error (MAE) of *Z*_*t*_ and CI width, respectively. Autocorrelation structure is autoregressive order of 4 (AR4).

For 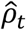, sFC cannot capture the dynamic changes and has a larger MAE compared to the other four dynamic methods, which are the same because they have the same *w*_1_ for 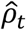 (Figure 2a). The MAE for dynamic methods is relatively high during the transition between the two stage pattern, since the sliding window method can capture continuous changes, rather than the stepwise changes. For 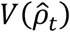 , overall, DSW method has the smallest mean MAE, compared to other methods, illustrating the advantage of considering dynamics in correlation, ACF and XCF with dual window size. In the second stage, SSW is most close to the truth, followed by DSW method. However, in the first stage, SSW has a very high variance estimation and large MAE (Figure 2b).

Figure 3a shows for CI coverage of true correlation, SSW and DSW are most close to 95%. Note that because of the stepwise change in true dFC in the simulation, the CI coverage of true correlation is small during the transition between the two stages (Figure 3a). We expect CI coverage of zero correlation increases as 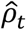 decreases from first stage to second stage. While other methods follows this expectation, the SSW has extremely high CI coverage of zero correlation (above 70%) at the first stage, suggesting the inference of insignificant correlation in more than 70% of simulation repetitions when the true correlation is 0.9 (Figure 3b). This is an egregious error and leads to faulty inference in terms of identifying dynamic connectivity. This is consistent with what we observed in Figure 2b, how the SSW substantially overestimates 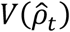 in the first stage.

For *Z*_*t*_, DSW is most close to the simulation truth and has the smallest mean MAE, compared with other methods. SSW has a smaller Z score estimation in the first stage, compared with other methods (Figure 4a). Looking at the CI width results (Figure 4b), we expect CI width increases as 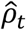 decreases from first stage to second stage, since 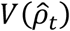 increases as 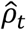 decreases. Note that the truth of CI width is relatively high during the transition between stages due to simulation artifact. However, SSW does not follow this expectation as other methods do, with a larger CI width at the first stage. This is consistent with what we observed in Figure 2b and Figure 4a, how the SSW substantially overestimates 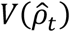 , and consequently underestimates *Z*_*t*_ at the first stage of high correlation values. This kind of poor CI behavior is because the static XCF in SSW cannot capture the highly dynamic XCF when 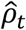 is large. Overall, considering the poor CI performance of SSW, we conclude that under this scenario, DSW has the best overall performance.

To further illustrate the CI and the compare DSW and SSW, we take a closer look at a simulation pair of ROIs and in vivo resting-state fMRI data example (Figure 5). Figure 5a shows for a simulation pair of ROIs under the above two stage AR4 scenario, DSW has narrower CIs that follows the dynamic correlation. However, SSW has very large CIs at the first stage when correlation is very high, with some intervals including zero correlation, altering the statistical inference. For Z scores, SSW cannot capture the relatively high Z scores at the first stage, while the Naïve overestimates Z scores. DSW can capture the dynamics and mitigate the overestimation, consistent with the simulation results in Figure 4. Similar results can be observed for the in vivo resting-state fMRI data example, same example as in Figure 1a (Figure 5b). SSW has wider CIs, especially at around scan 180 and 400, altering the direction of statistical inference, consistent with Z score plots.

**Figure 5.**
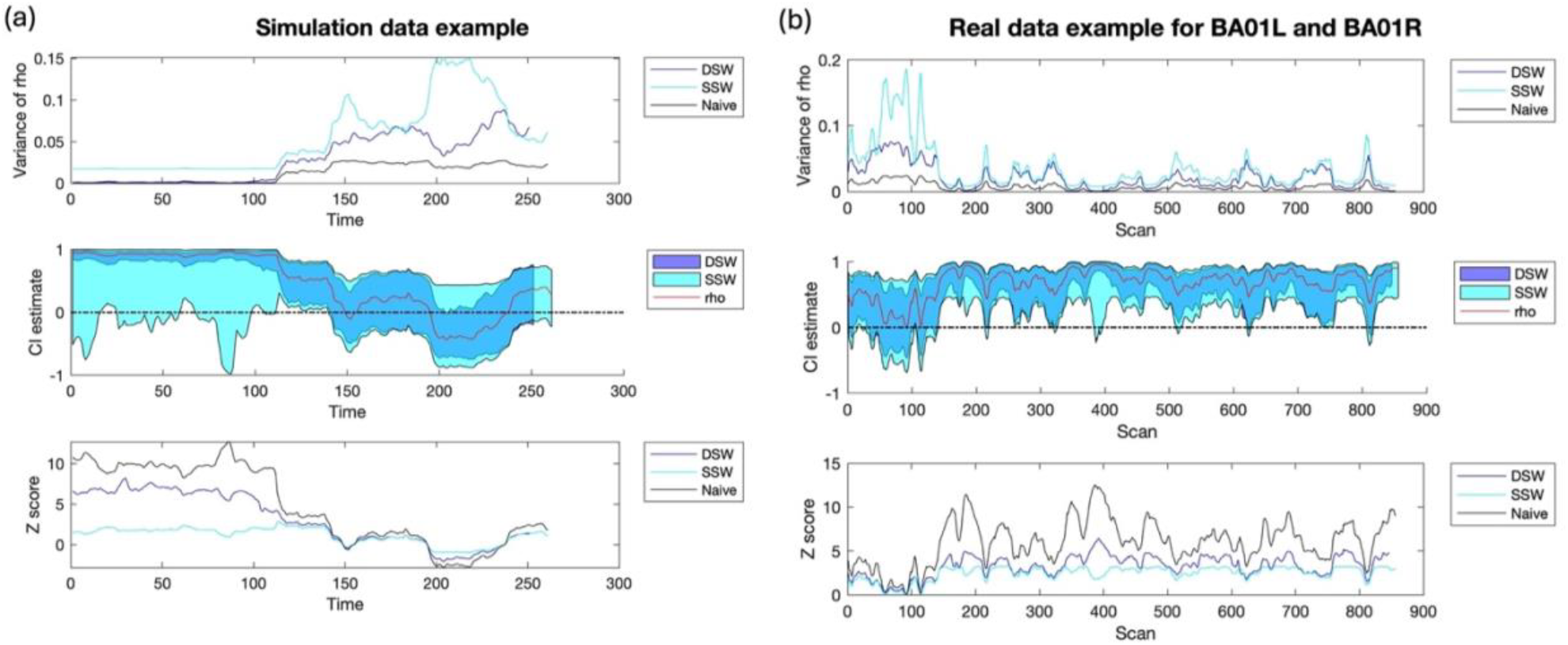
Pair of ROIs examples comparing different variance estimators. (a) simulation data example. (b) In vivo data example for Brodmann Area 1 at left (BA01L) and Brodmann Area 1 at right (BA01R).

Simulation results for another scenario, Pattern 3 monotonic decreasing with AR4 structure, are shown in Supplement Figure 3. For 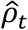 , sFC has a larger MAE compared to the other four dynamic methods (Supplement Figure 3a). For 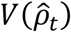 , SSW and DSW are most close to the simulation truth and has the smallest mean MAE (Supplement Figure 3b). Similar results is observed for *Z*_*t*_ (Supplement Figure 3c). However, SSW substantially overestimates 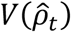 and underestimates *Z*_*t*_ at sliding window 1-50, where correlation is relatively high in the dynamics. For the CI results (Supplement Figure 3d-e), SSW has a large CI width and CI coverage of zero at sliding window 1-50, when 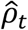 is 0.8-0.9. For CI coverage of true correlation, SSW and DSW are closest to 0.95. Thus, consistent with the previous scenario, we conclude that DSW has the best overall performance.

A summary of simulation results for all scenarios is shown in Figure 6. DSW and SSW has the smallest mean MAE for variance, compared with other methods. However, since SSW has poor CI performance when the correlation is relatively high during the a portion of the scans, possibly altering the statistical inference, we conclude from that DSW is a better variance estimator compared to others.

**Figure 6.**
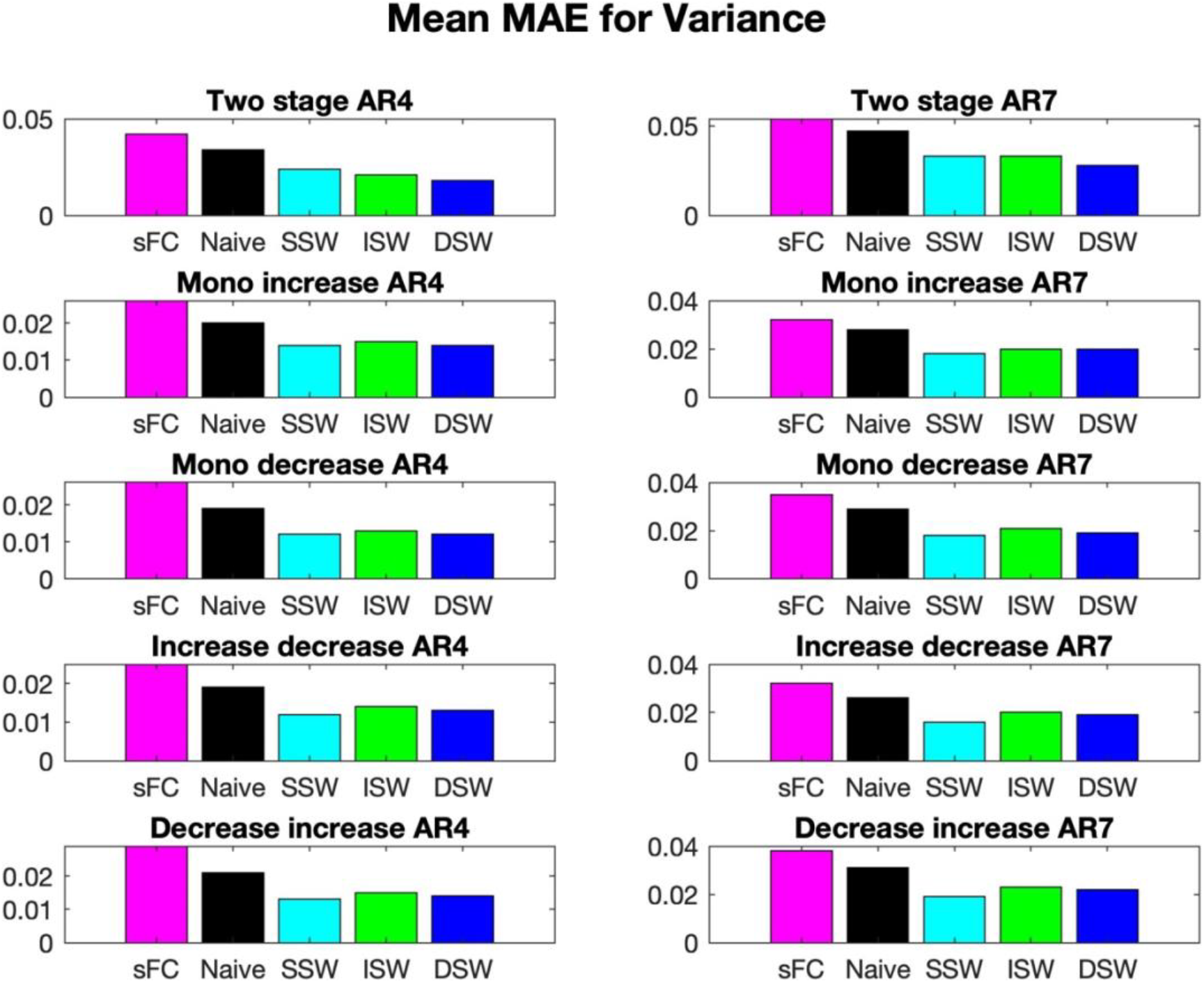
Summary of simulation results for all scenarios. Mean of Mean Absolute Error (MAE) across repetitions and windows. AR4: Autocorrelation structure with order of 4. AR7: Autocorrelation structure with order of 7.

### 3.2 Window size recommendation

Table 1 shows the results for window size selection with a medium tolerance percentage *q* as 10%, permitting an allowable 10% increase in MAE from the minimum mean MAE. The window size for XCF is 1.33 (1.23, 1.38) (median (1^st^ Quantile, 3^rd^ Quantile)) times the window size for correlation. An example for window size and MAE for a simulation scenario is shown is Supplement Figures 4. For a small tolerance percentage *q* as 5%, the window size for XCF is 1.50 (1.21, 1.67) times the window size for correlation (Supplement Tables 1). For a large tolerance percentage *q* as 15%, the window size for XCF is 1.50 (1.28, 1.60) times the window size for correlation (Supplement Tables 2). Note that without tolerance percentage (*q* = 0), the window size for XCF is 1.50 (1.31, 1.50) times the window size for correlation. The results are consistent between different tolerance percentages. Thus, we suggest to choose a window size for ACF and XCF that is 1.2-1.5 times the window size for correlation. As a conservative approach, if a ratio is to be applied across the whole brain, we suggest to apply 1.2 times. This is to insure that the unwelcome CI phenomena as suffered by SSW does not arise through overly long windows for some of the pairs of ROIs.

**Table 1.**
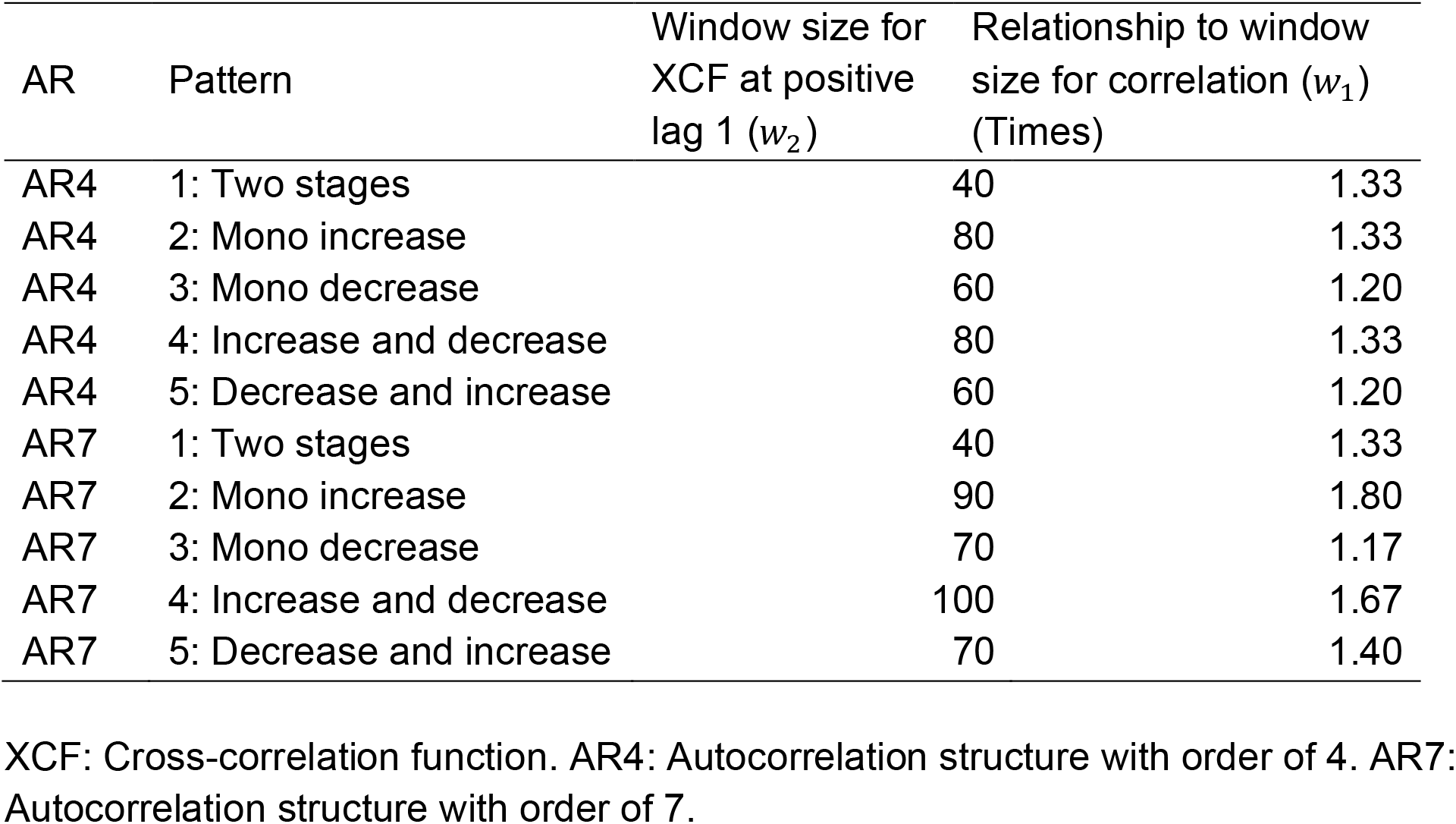
Window size selection for different scenarios, tolerance = 10%.

### 3.3 In vivo resting-state fMRI data results

The mean of sFC rho across subjects is consistent with previous study (Vakamudi et al., 2020) and slightly larger than the mean of dFC rho across windows and subjects (difference=0.045, *p*<0.001) (Supplement Figure 5a, 5b). The STD (Standard Deviation) of dFC across windows and subjects is significantly larger than the STD of sFC across subjects (difference=0.160, *p*<0.001) (Supplement Figure 5c, 5d), demonstrating significant variation of dFC rho across windows. The means of sFC Z scores across subjects are larger than the means of dFC Z scores across windows and subjects (difference= 5.014, *p*<0.001) (Supplement Figure 6a, 6b), because of smaller variance for sFC rho due to more data being available. STD of sFC Z scores across subjects are larger than STD of dFC Z scores across windows and subjects (difference=0.8176, *p*<0.001) (Supplement Figure 6c, 6d), since sFC Z scores have a larger mean. The mean of Coefficients of Variation (CV) of sFC Z scores across subjects are smaller than the mean of CV of dFC Z scores across windows and subjects (difference= 0.351, *p*<0.001) (Supplement Figure 6e, 6f), consistent with the comparison between STD of sFC and dFC, demonstrating significant variation of dFC across windows.

Besides the mean of dFC Z scores, our method provides more information about the dFC in the form of percentage of dFC CI that covers zero correlation (CI coverage of zero correlation) and percentage of dFC CI that covers sFC (CI coverage of sFC). They demonstrate the significance of the dFC correlations and the level of dynamics, respectively. CI coverage of zero correlation, serving as a summary statistic for uncertainty estimation in dFC, indicates whether the dFC for a pair of ROIs is predominantly significant, predominantly insignificant, or a mixture of both significant and insignificant connectivity, with values close to 0%, 100%, and 50%, respectively. The mean of CI coverage of zero correlation across subjects informs similar network information as the mean of sFC rho, showing most significant dFC in dark blue color and least significant dFC in dark red color (Figure 7a). Mean of CI coverage of sFC across subjects shows how dynamics each pair of ROis is, with larger percentages representing less dynamic connectivity and smaller percentages representing more dynamic connectivity (Figure 7c). The STD of CI coverage of zero correlation and sFC across subjects shows the variation between subjects (Figure 7b, 7d).

**Figure 7.**
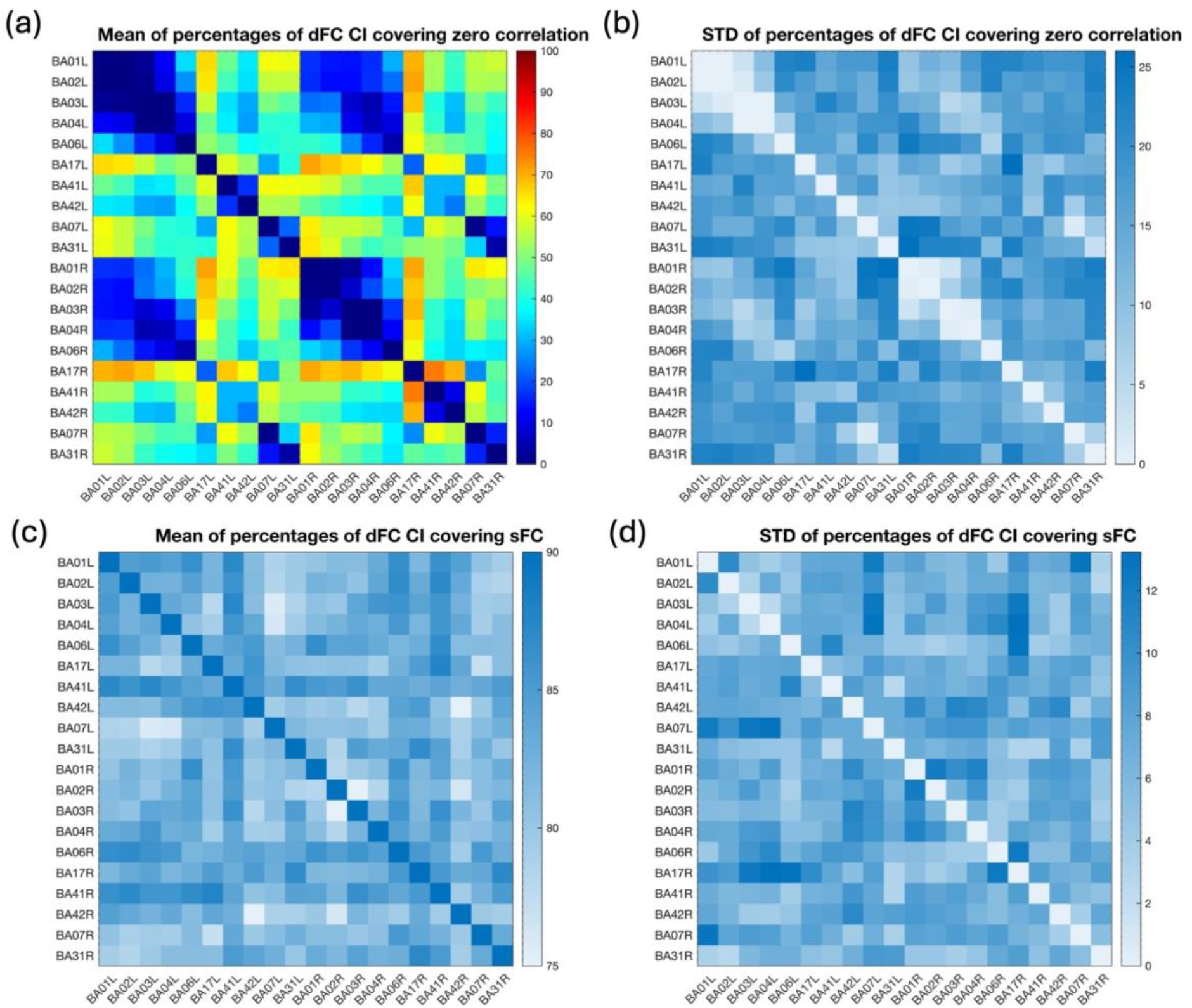
Mean and standard deviation (STD) of percentages of dynamic Functional Connectivity (dFC) Confidence interval (CI) covering zero correlation (CI coverage of zero correlation) and static functional connectivity (sFC) (CI coverage of sFC) across subjects. Dual Sliding Window (DSW) variance estimator for sliding window method is used with *w*_1_ =45 (18 seconds) and *w*_2_=55 scans (22 seconds)

Additionally, using simulations, we benchmark the distribution of CI coverage of sFC for a pair of ROIs that has sFC as truth (Supplement Figure 7). With the truth of sFC varying from 0.2 to 0.8, median of CI coverage of sFC is consistently around 90%-91% for AR4 and 87%-88% for AR7. These values are higher than most of the percentages observed in Figure 7c. This benchmarking result further demonstrates that, for most pairs of ROIs in Figure 7c, the truth of sFC does not hold, indicating that most pairs of ROIs exhibit dynamic behavior. With the code provided on GitHub, researchers can customize the benchmarking of the distribution of CI coverage of sFC by specifying parameters such as the sFC correlation value, AR structure, and the total number of scans. This allows researchers to determine the threshold of CI coverage of sFC for a given pair of ROIs with sFC by simulation. If the percentage is smaller than this threshold, it can be concluded that the pair of ROIs exhibits dynamic behavior.

Zooming into pair of ROI results, the dFC between BA01L and BA02L is very significant with small variation between subjects (CI coverage of zero correlation 0% for all subjects), while the dynamics varies between subjects (CI coverage of sFC 94.1%, 92.1%, 81.0%, 83.8%, 91.1% and 65.7% for subject 1-6) (Figure 8a), indicating significant dFC correlation despite highly dynamic dFC. The dFC between BA42L and BA42R is highly correlated with large variation between subjects (CI coverage of zero correlation 41.6%, 3.4%, 44.9%, 3.2%, 38.1% and 15.7% for subject 1-6), and the dFC is highly dynamic with large variation between subjects (CI coverage of sFC 85.8%, 80.7%, 59.1%, 63.0%, 80.7%, 73.7% for subject 1-6). The dFC between BA01L and BA06L is moderately correlated with high variation between subjects (CI coverage of zero correlation 26.8%, 18.1%, 6.5%, 36.1%, 48.5% and 68.6% for subject 1-6), and the dFC is less dynamic with small variation between subjects (CI coverage of sFC 82.6%, 82.5%, 91.4%, 83.9%, 93.9%, and 83.92% for subject 1-6). This can be observed in subject level plots too (Supplement Figure 8). Compared to the sFC correlation and Z scores, more information regarding statistical significance and dynamics can be obtained using our method.

**Figure 8.**
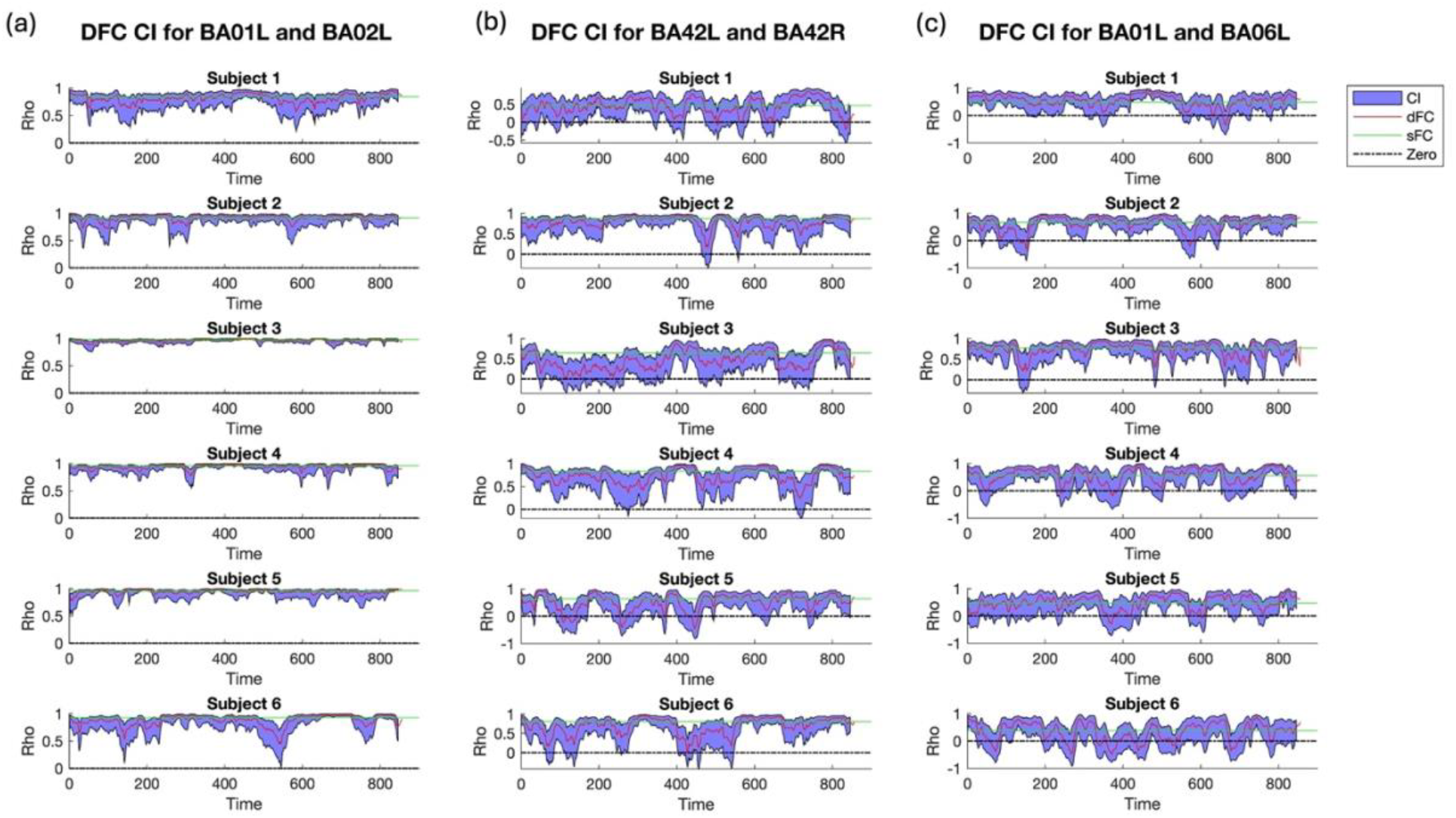
Dynamic functional connectivity (dFC) Confidence interval (CI) for three pairs of ROIs for each subject. (a) left Brodmann area 01 (BA01L) and left Brodmann area 02 (BA02L), (b) left Brodmann area 42 (BA42L) and right Brodmann area 42 (BA42R), (c) left BA01L and left Brodmann area 06 (BA06L). Dual Sliding Window (DSW) variance estimator for sliding window method is used with *w*_1_ =45 (18 seconds) and *w*_2_=55 scans (22 seconds)

The advantages of DSW compared with SSW with in vivo resting-state fMRI data are shown in Figure 9. Mean of CI coverage of sFC derived with SSW variance estimator has a median of 98.7% across pairs of ROIs, and is significantly larger than with DSW variance estimator (difference = 16.34%, *p*<0.001). It demonstrates that SSW, which does not considers the dynamics in ACF and XCF, cannot capture the dynamics in functional connectivity, consistent with simulation results. CI for BA42L and BA42L (Figure 9b) shows wider CI compared to results in DSW (Figure 8b) and poor CI behavior during the high correlation stage of the dynamics, such as Subject 5 at around scan 50 and 200.

**Figure 9.**
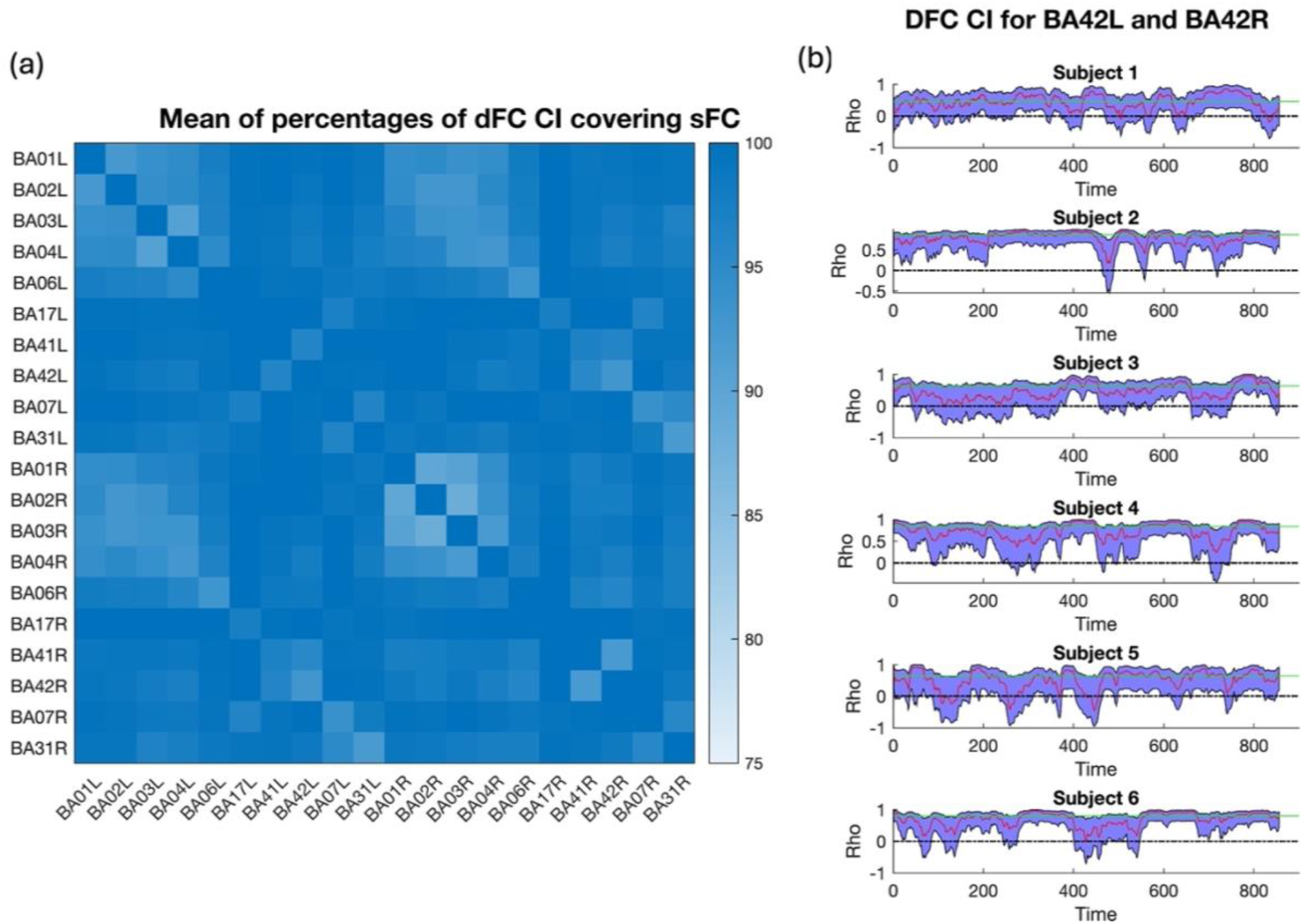
Statistics using Static ACF and XCF Sliding Window (SSW) variance estimator with window size 45 scans (18 seconds) (a) Mean of percentages that dynamic Functional Connectivity (dFC) Confidence interval (CI) covers static functional connectivity (sFC) (CI coverage of sFC) across subjects (b) dFC CI between left Brodmann area 42 (BA42L) and right Brodmann area 42 (BA42R) in each subject.

We also examined DSW results with *w*_1_ =75 (30 seconds) and *w*_2_ =90 (36 seconds) (Supplement Figure 9-11). The results are consistent with DSW results with *w*_1_ =45 (18 seconds) and *w*_2_=55 scans (22 seconds). The framewise displacements plot for every patient indicates small magnitude of random movement that is not related to the observed dynamics (Supplement Figure 2).

## 4. Discussion

The performance of sliding window correlations has been questioned, mainly because it is challenging to decide if the fluctuations in an observed dFC time series are due to statistical uncertainty or reflect true dFC changes (Hindriks et al., 2016; Leonardi and Van De Ville, 2015; Shakil et al., 2016). Our method provides insights into this issue by quantifying the confidence intervals for the sliding window correlations (Preti et al., 2017; Solo et al., 2018). By examining the CI coverage of zero correlation across the scan, we can determine whether the dFC for a pair of ROIs is predominantly significant, predominantly insignificant, or a mixture of both significant and insignificant connections. Looking at the CI coverage of sFC across the scan, we can assess whether the connection for a pair of ROIs is more dynamic or more static. By reviewing the mean and standard deviation of Z scores across the scan, we can gain insight into the magnitude of the connection and the variation of the dynamics.

From the simulation results, we see that the Naïve variance estimator does not consider ACF and XCF and underestimates variance and overestimates Z score values.

Considering autocorrelation is crucial due to its significant impact on the inference of dFC (Honari et al., 2019). The SSW variance estimator only considers the static ACF and static XCF, ignoring the dynamics in ACF and XCF, which can lead to poor CI behavior and underestimated Z scores when correlation is relatively large during a portion of the scans. ISW uses shared window sizes for the estimation of correlation and covariance (ACF and XCF) and underperforms compared to DSW. Our DSW variance estimator not only considers the dynamics in ACF and XCF, but also allows dual window sizes for correlation, ACF and XCF for better estimation. This approach is the best estimation approach.

Based on the in vivo resting-state fMRI data analysis results, we see that our method can give information beyond a single sFC correlation value or a series of dFC sliding window correlation values. With the DSW variance estimator for sliding window correlations, more information about the dFC, such as the percentage of dFC CI that covers zero correlation and variability of dFC Z scores, can be obtained. Aiming at the application in preoperative and intra-operative fMRI for brain tumor resection, neurosurgeons can make informed decisions with information such as the percentage of dFC CI that covers zero correlation about the eloquence of brain tissue regions near the tumor. The addition of intraoperative fMRI to intraoperative structural MRI thus has the potential to improve intra-operative surgical guidance by co-localizing functional information and anatomical structures.

The ACF and XCF estimation is challenging in time series data. In our method, we use the Fast Fourier Transform (FFT) method based on the Wiener–Khinchin theorem, which does not assume a specific autoregressive order and can estimate all lags computationally efficiently. We regularize the ACF and XCF using Tukey Tapering method, since the ACF and XCF are expected to diminish to zero with increasing lags. We show that with limited sliding window data, ACF and XCF estimates from FFT method can decently approximate the true parameters using simulations (Supplement Figures 12-14).

We observe significant and consistent temporal fluctuations in ACF and XCF across lags in multiple pairs of ROIs and subjects. The high sensitivity of the high-speed fMRI (TR 0.4 seconds) employed in this study help the detection of these dynamic fluctuations in ACF and XCF, allowing us to capture subtle and rapid changes in neural activity. Besides the direct estimation of dynamics in ACF and XCF, the poor performance of SSW also highlights the importance of considering the temporal dynamic fluctuations in ACF and XCF.

In our simulations, we use the step-wise function for dynamic patterns of rho, because of simulation limitations. In practice, the resting state dynamic correlation tends to change gradually (Kudela et al., 2017) and we expect better performance comparied to the simulation results. The simulation truth for the variance of sliding window correlations is generated via Monte Carlo simulations (Afyouni et al., 2019). Because the theoretical variance of Pearson correlation coefficient with autocorrelated data cannot be directly calculated using traditional formulas. Thus, we use the Monte Carlo simulation truth as the ground truth for different methods to compare with.

We noticed that compared to the simulated true variance, all variance estimators tend to underestimate the truth. This is because the FFT method for ACF and XCF estimation tends to underestimate the parameters with shorter time series. We attempt to alleviate this problem by using a slightly larger window size for the ACF and XCF estimation, which appears helpful since the DSW outperforms the ISW. However, like the window size for sliding window correlations, the window size for ACF and XCF cannot be too large or the dynamics would not be averaged out, as seen with SSW performance.

The window size selection for sliding window correlation is not a trivial question and has been studies by many researchers (Savva et al., 2019). There are several considerations when selecting the window size. First, the temporal dynamics of resting-state networks need to be considered. Typical choices of window size are 30 – 60 seconds, which are suggested to capture the cognitive state change based on empirical studies and method research (Leonardi and Van De Ville, 2015). The number of time points for the sliding window will depend on the Repetition Time (TR) of the real data.

Second, statistical consideration dictate a trade-off between bias and variance in the sliding window method. We use *w*_1_ for correlation as an example. Supplement Figure 15 shows in simulations, with a smaller window (eg., *w*_1_ = 30), the bias of the estimated correlation is small, but the variance is relatively larger. However, with a larger window (eg., *w*_1_ = 90), the bias is large, and the variance is relatively smaller. Thus, a balanced window that optimally balances the bias-variance trade-off is ideal.

Another consideration is from the information gain, that a larger window will result in a shorter series of available data for estimations of correlation. For example, in Supplement Figure 15, *w*_1_ = 30 can detect up to a correlation of 0.9 at window number 271, while *w*_1_ = 90 can only detect up to a correlation of 0.8 at window number 211, resulting in some loss of information. Thus, we do not want to increase window size for a relatively small decrease in MAE, which can be balanced using the rate of change (derivative) for MAE per unit window increase in our simulation. Our adopted approach leads to a similar range of window sizes as suggested by others. With our method, simulations suggest a rule-of-thumb approximate relationship between window sizes for *w*_1_ and *w*_2_ .

When considering multiple pairs of connections in the fMRI study, the correction for multiple comparison is important. The standardized Z scores can be corrected similarly as the p values using the False Discovery Rate. For the CIs of dynamic correlations, a conservative approach of Bonferroni correction can be used when looking at CI coverage of zeros or sFC.

In future optimization of our method, we will consider deciding between static and dynamic estimates of connectivity, depending on the data structure. In our simulations, we observe that if a connection is static, the best estimators are using static correlation and covariance models, as one would expect. In addition, we want to further use our method to dynamically adjust window sizes for each pair of ROIs.

## 5. Conclusion

Our proposed DSW variance estimator for sliding window correlation considers the dynamics in ACF and XCF. It is more consistently accurate in terms of statistical inference compared with stationary covariance model and single window model. It can provide statistics, such as dynamic Z scores, CI, percentages of dFC CI covering zero correlation and sFC, to dynamically characterize fluctuations of connectivity. This method can serve as a basis for determining between dynamic and static connectivity between pairs of ROIs.

