## Supplement for "Sliding window functional connectivity inference with nonstationary autocorrelations and cross-correlations"

### Supplement Materials

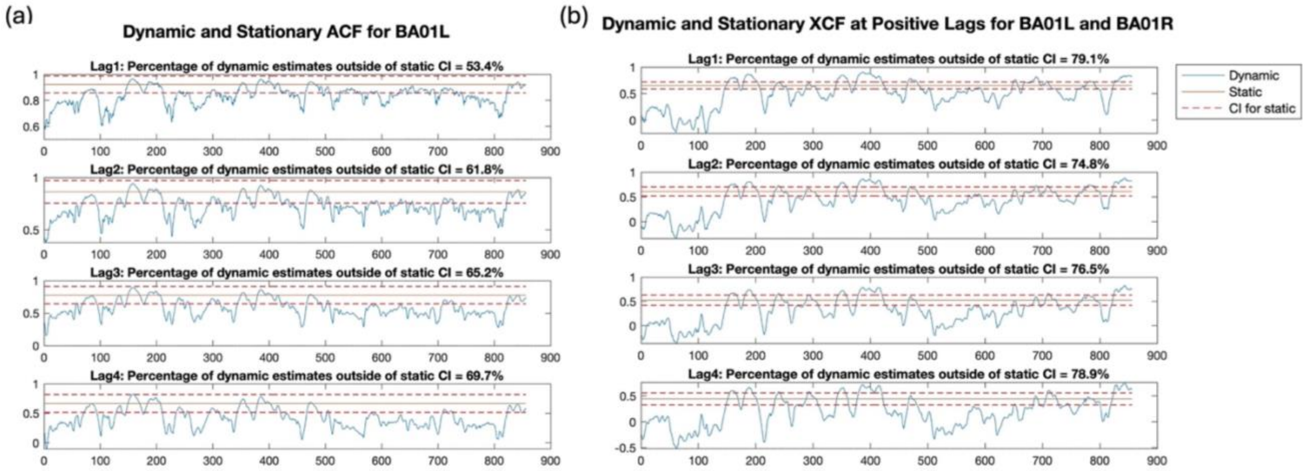


**Supplement Figure 1.** Dynamic Autocorrelation Functions (ACF) and Cross-correlation Functions (XCF) compared with 95% Confidence Interval (CI) for static ACF and XCF. The percentage of dynamic estimates outside of static CI is calculated along the time. (a) Dynamic ACF and 95% CI for static ACF for Brodmann Area 1 at left (BA01L) at lags 1-4. (b) Dynamic XCF and 95% CI for static XCF between BA01L and Brodmann Area 1 at right (BA01R) at positive lags 1-4

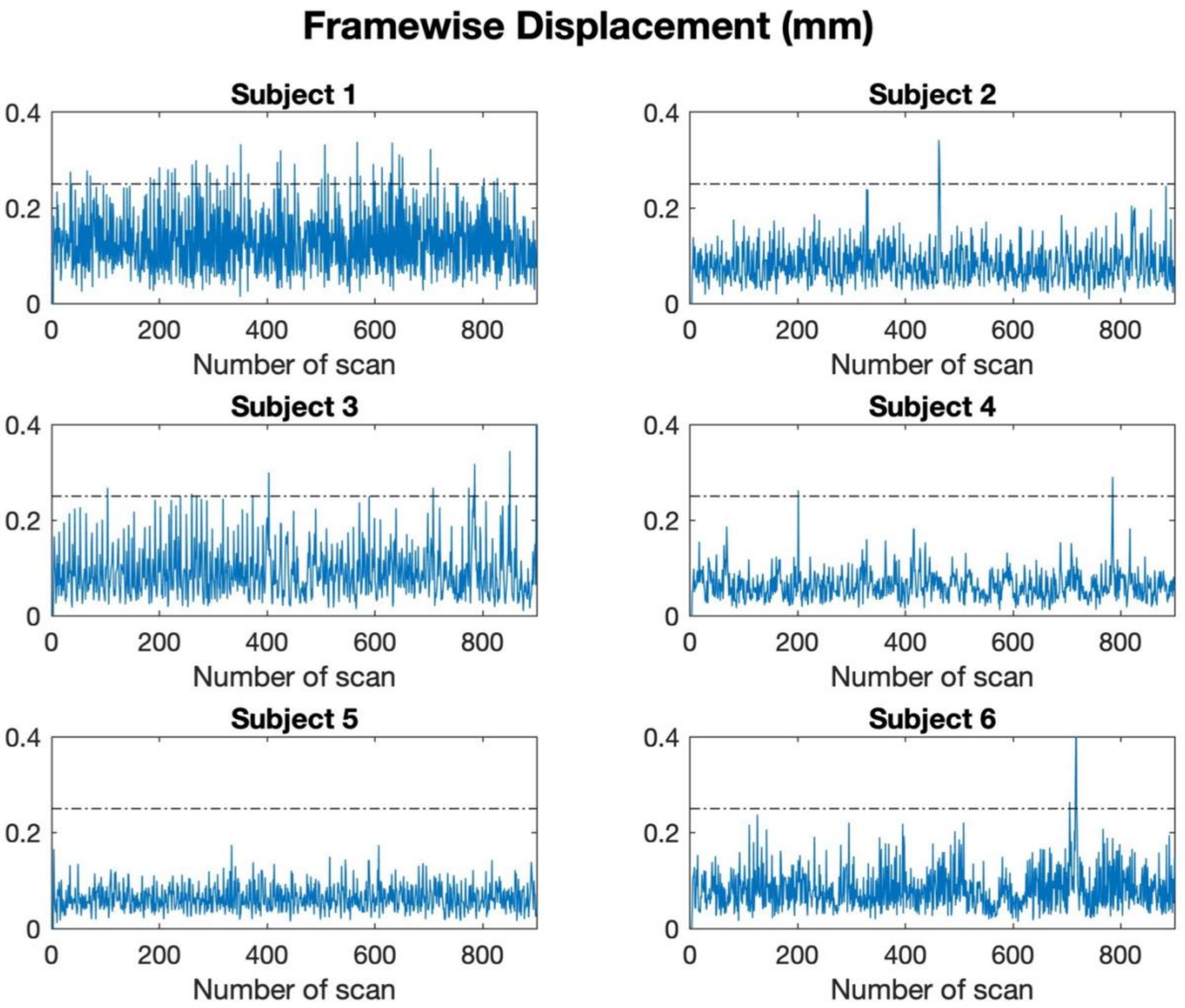


**Supplement Figure 2.** Framewise displacement in millimeters for each subject

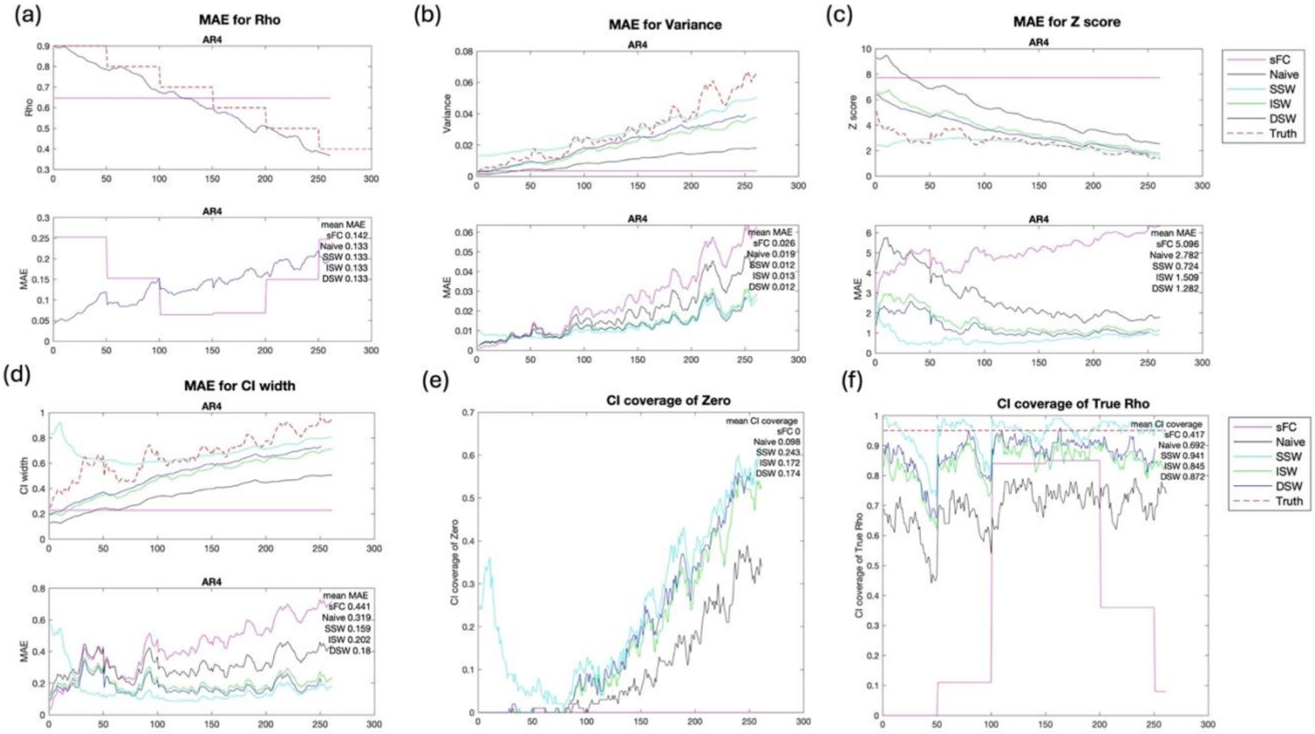


**Supplement Figure 3**. Simulation results comparing different variance estimators for simulation scenario with dynamic correlation Pattern 3 (monotonic decreasing) with AR4 structure. (a)-(d) Upper panels in each figure are the mean of $\hat{\rho}_t$, $V(\hat{\rho}_t)$, $Z_t$ and Confidence Interval (CI) width, respectively, among 100 simulations for different methods. Lower panels in each figure are Mean Absolue Error (MAE) of $\hat{\rho}_t$, $V(\hat{\rho}_t)$, $Z_t$ and CI width, respectively. AR4: Autocorrelation structure with order of 4.

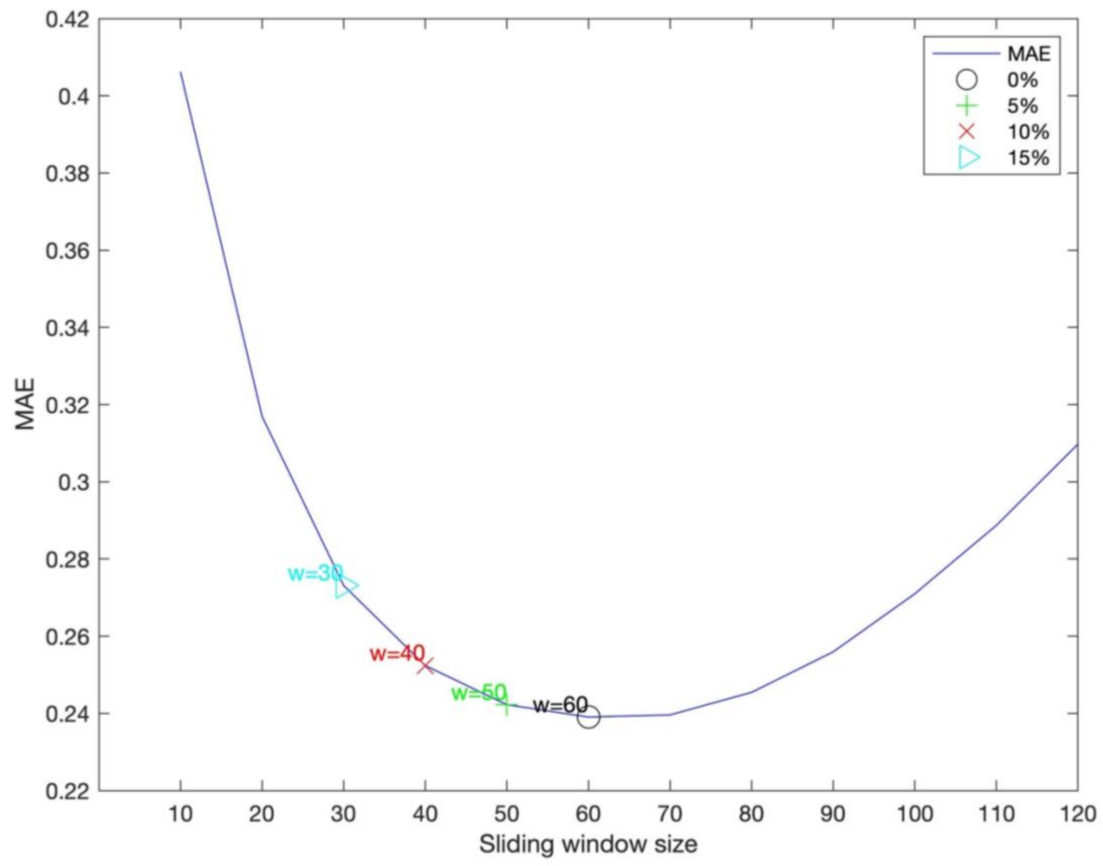


**Supplement Figure 4.** An example of window size selection for cross-correlation (XCF) at positive lag 1 using Mean Absolute Error (MAE) and tolerance percentages. This scenario is Two stage with AR7 structure and total time points 300. Black circle is the window size without tolerance percentages. The window sizes with tolerance percentages (5%, 10%, and 15%) are represented using green, red and cyan, respectively.

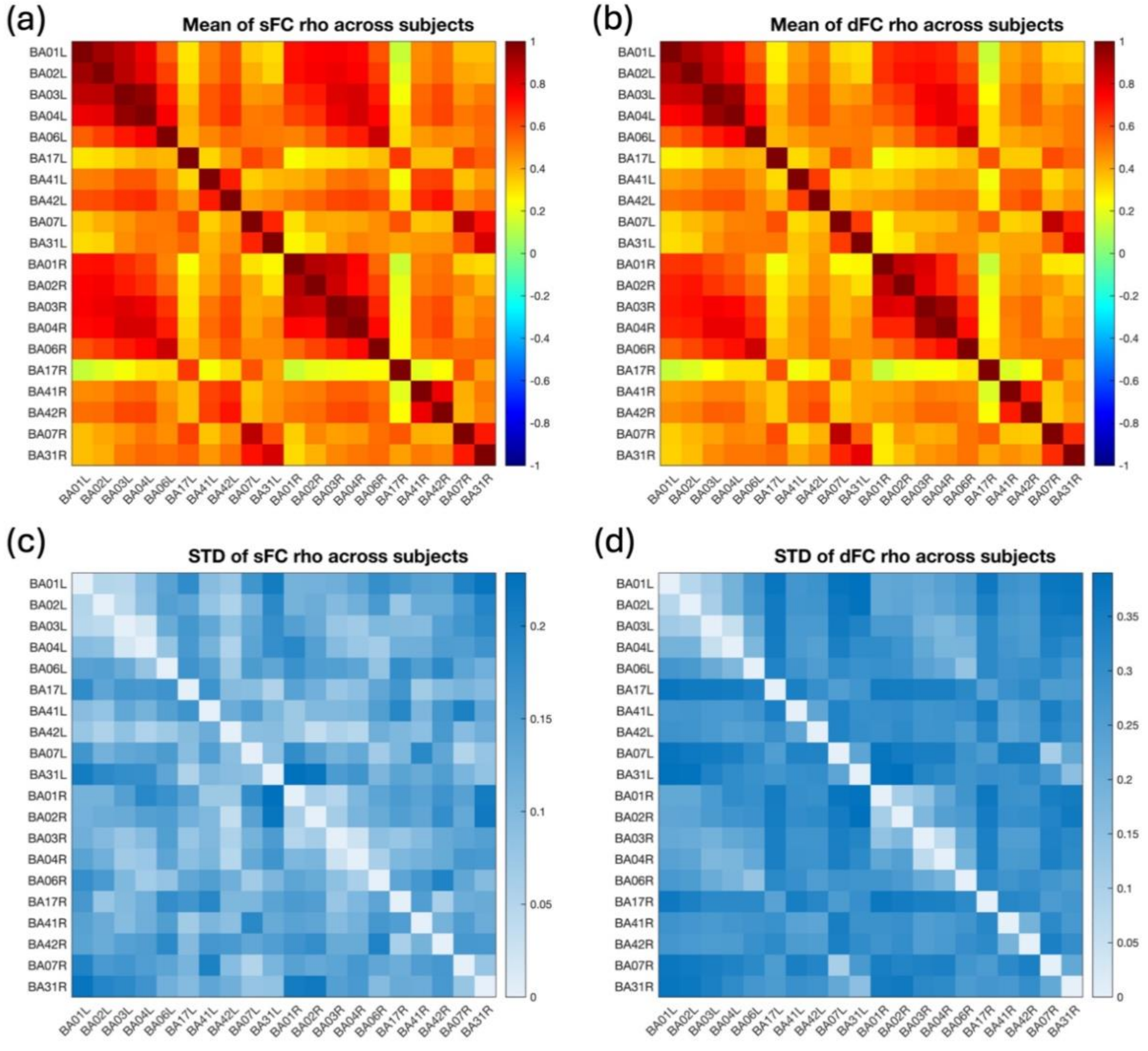


**Supplement Figure 5.** Mean and standard deviation (STD) of static functional connectivity (sFC) rho across subjects and dynamic functional connectivity (dFC) rho across windows and subjects

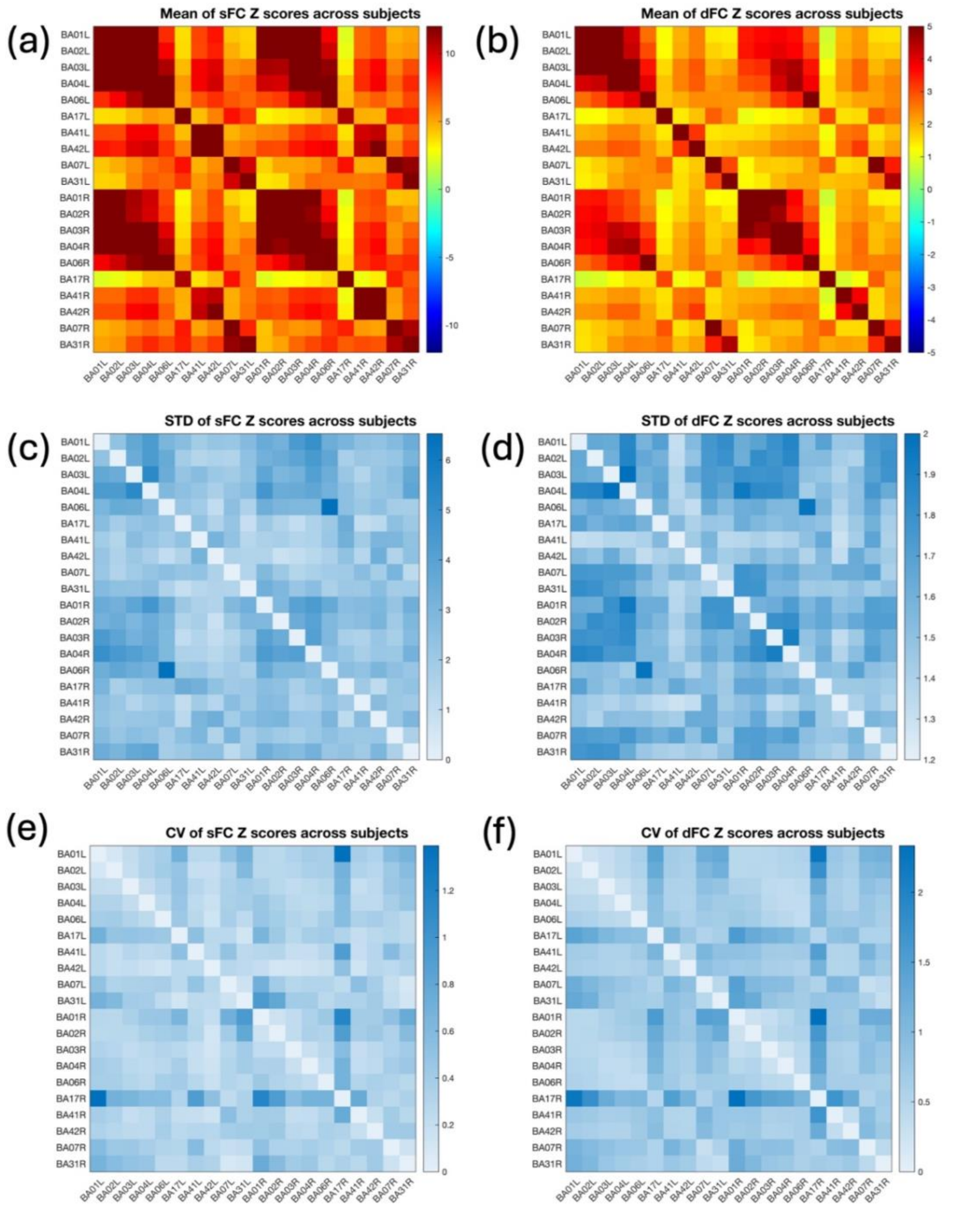


**Supplement Figure 6.** Mean, standard deviation (STD), and coefficient of variation (CV) of static functional connectivity (sFC) Z scores across subjects and dynamic functional connectivity (dFC) Z scores across windows and subjects.

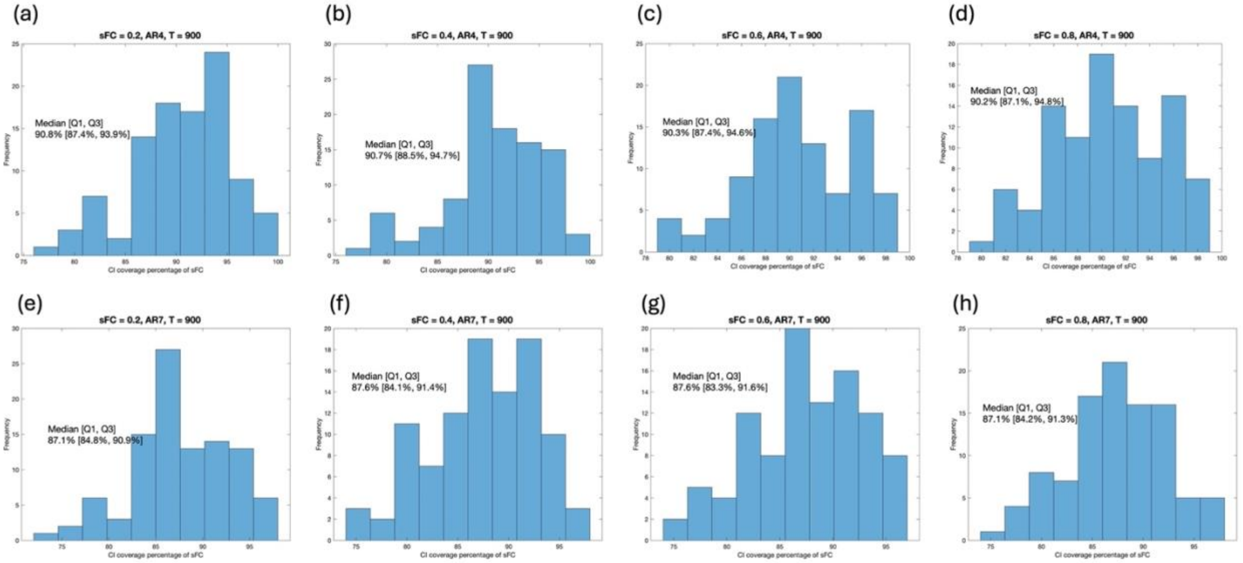


**Supplement Figure 7.** Histograms of the percentages of dFC CI that covers sFC for a pair of ROIs that has sFC as truth. Dual Sliding Window (DSW) variance estimator for sliding window method is used with $w_1$ =45 (18 seconds) and $w_2$=55 scans (22 seconds). dFC: dynamic Functional Connectivity. CI: Confidence Interval. sFC: static Functional connectivity. ROIs: Regions of Interest. AR4: Autocorrelation structure of order 4. AR7: Autocorrelation structure of order 7. T: total number of scans per ROI

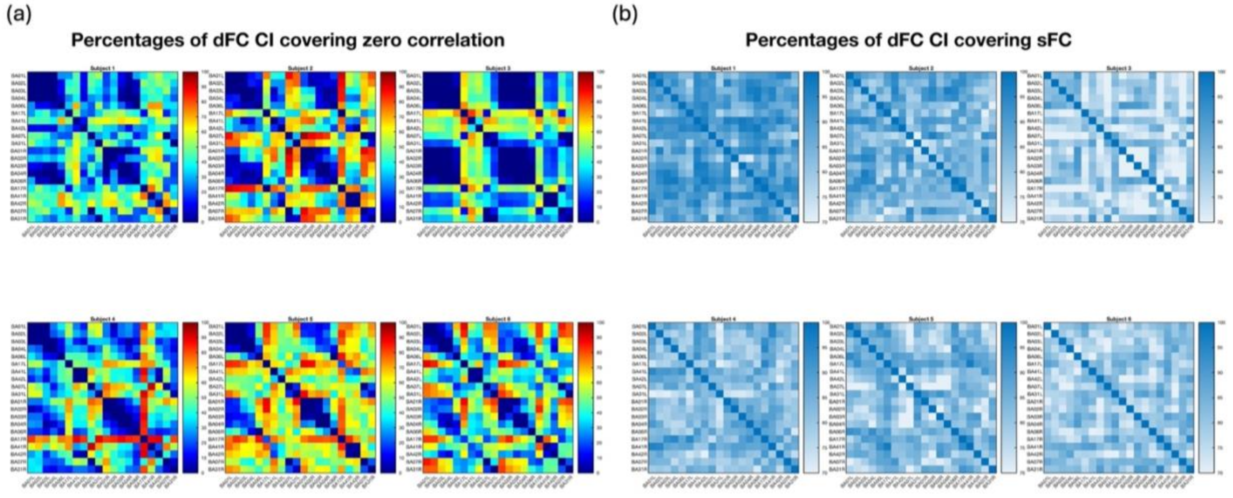


**Supplement Figure 8.** Percentages of dynamic Functional Connectivity (dFC) Confidence interval (CI) covers zero correlation and static functional connectivity (sFC) for each subject.

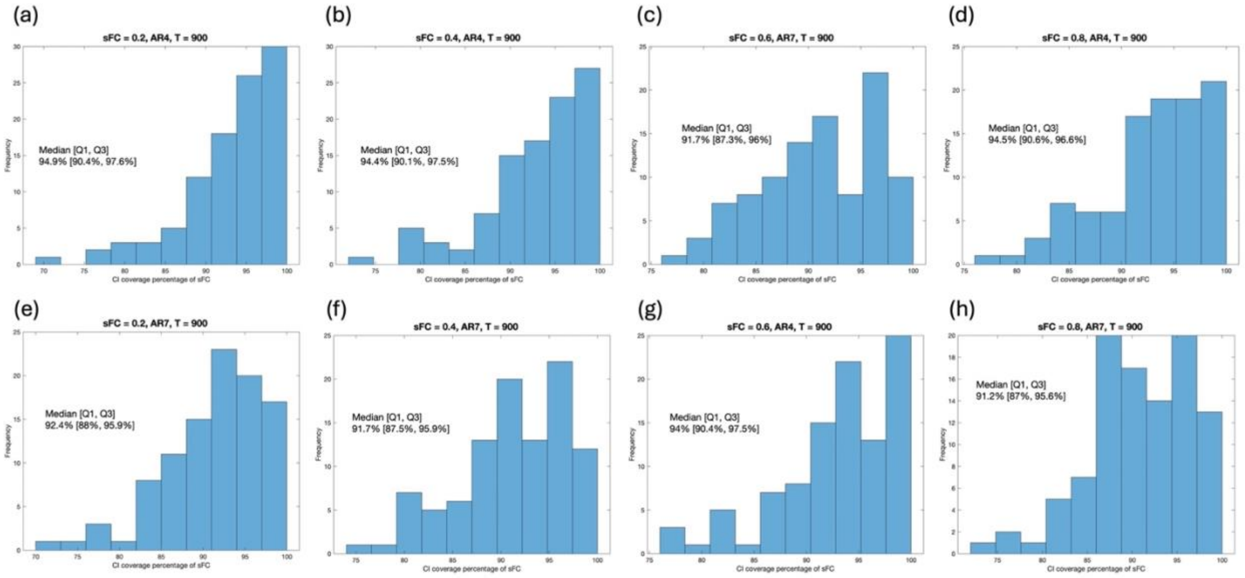


**Supplement Figure 9.** Histograms of the percentages of dFC CI that covers sFC for a pair of ROIs that has sFC as truth. Dual Sliding Window (DSW) variance estimator for sliding window method is used with $w_1$ =75 (30 seconds) and $w_2$ =90 (36 seconds). dFC: dynamic Functional Connectivity. CI: Confidence Interval. sFC: static Functional connectivity. ROIs: Regions of Interest. AR4: Autocorrelation structure of order 4. AR7: Autocorrelation structure of order 7. T: total number of scans per ROI

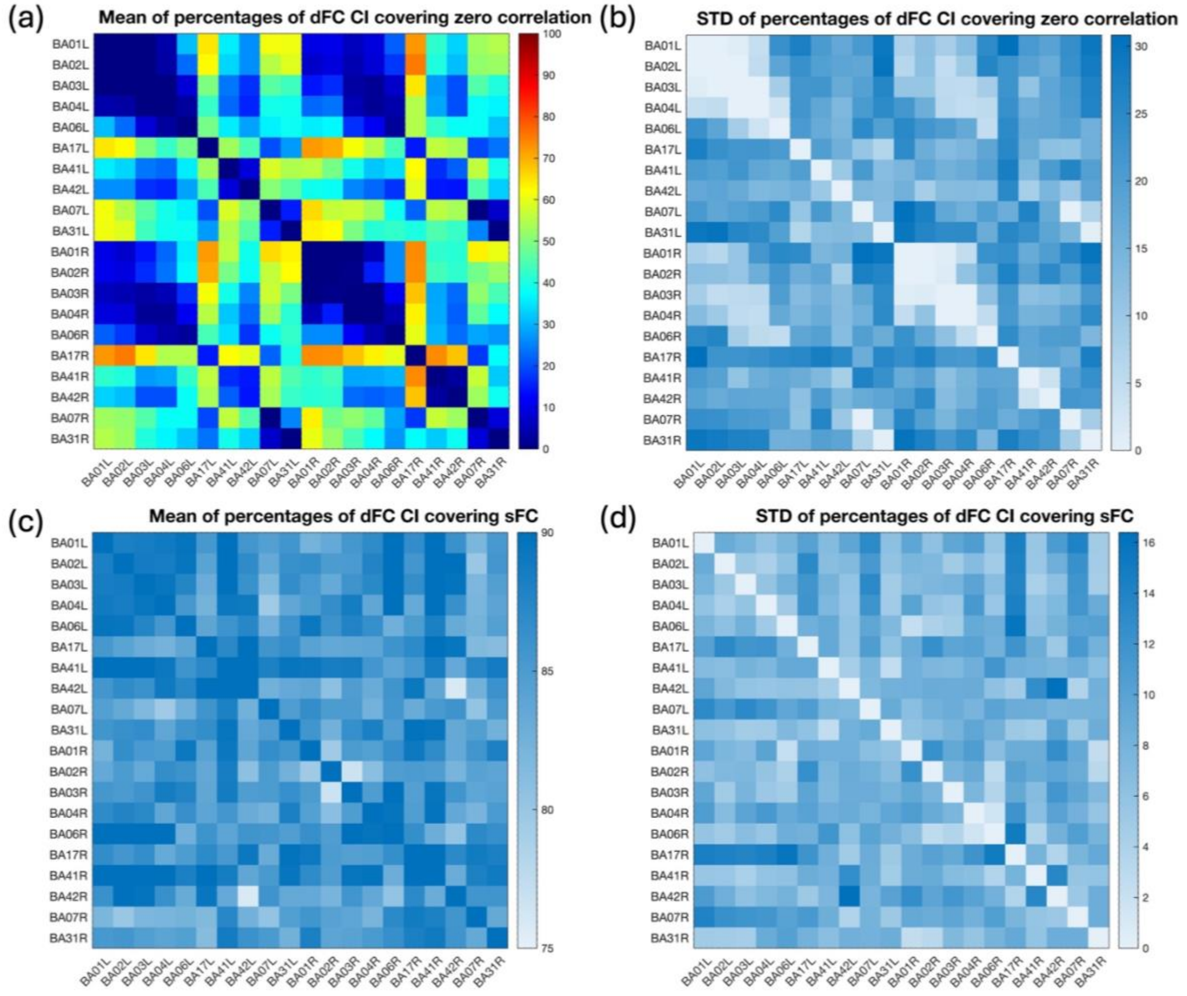


**Supplement Figure 10.** Mean and standard deviation (STD) of percentages that dynamic Functional Connectivity (dFC) Confidence interval (CI) covers zero correlation and static functional connectivity (sFC) across subjects. Dual Sliding Window (DSW) variance estimator for sliding window method is used with $w_1$ =75 (30 seconds) and $w_2$ =90 (36 seconds)

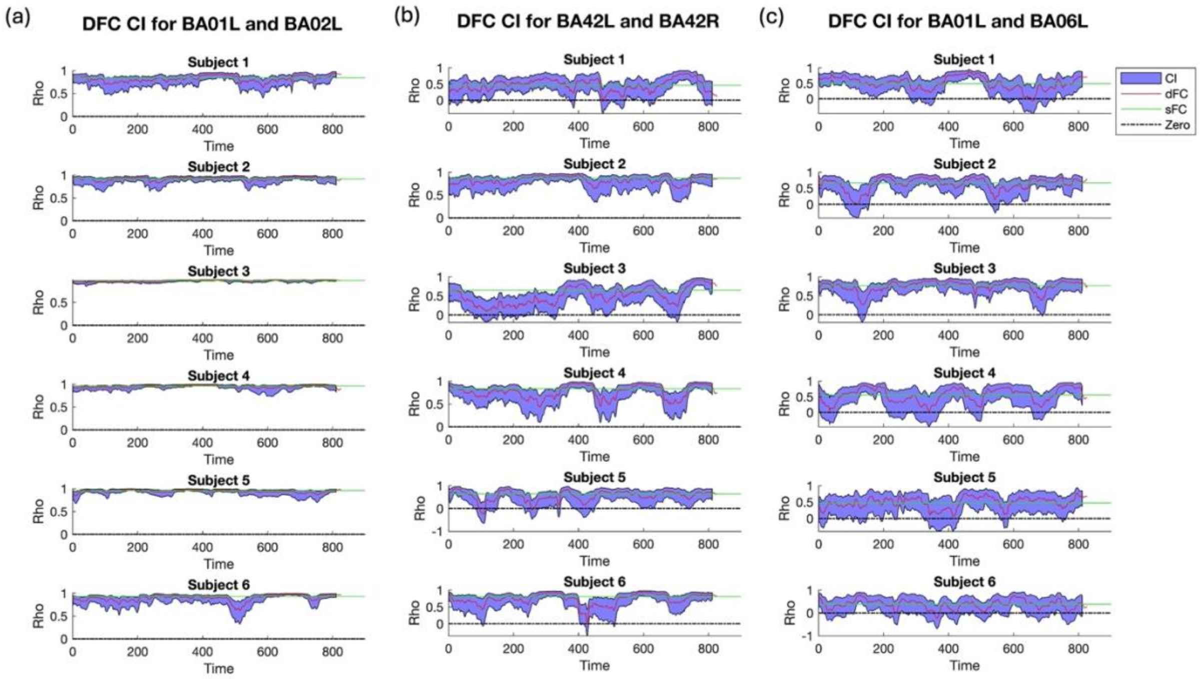


**Supplement Figure 11.** Dynamic functional connectivity (dFC) Confidence interval (CI) for three pairs and each subject. (a) Pair Brodmann Area 01 at Left side (BA01L) and Brodmann Area 02 at Left side (BA02L), (b) Pair Brodmann Area 42 at Left side (BA42L) and Brodmann Area 42 at Right side (BA42R), (c) Pair BA01L and Brodmann Area 06 at Left side (BA06L). Dual Sliding Window (DSW) variance estimator for sliding window method is used with $w_1$ =75 (30 seconds) and $w_2$ =90 (36 seconds)

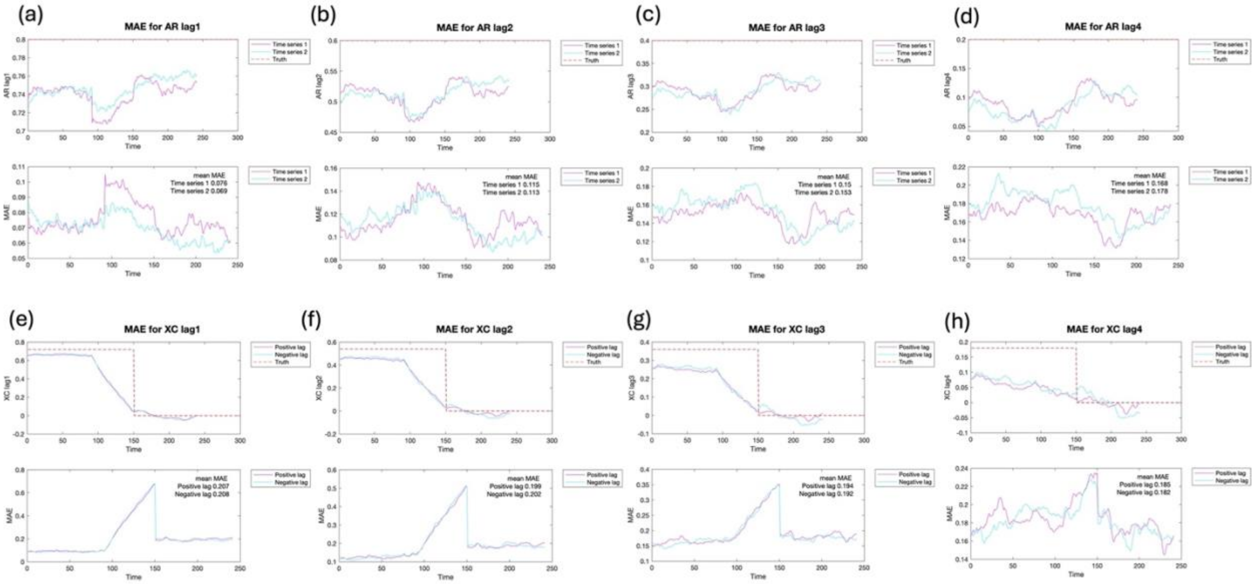


**Supplement Figure 12.** Simulation results of the autocorrelation function (ACF) and cross-correlation function (XCF) estimation using the Fast Fourier Transform (FFT) method. This simulation scenario is Pattern 1 (two stage) for dynamic correlation with AR4 structure. (a)-(d) ACF at lag 1-4. (e-h) XCF at lag 1-4. Upper panels in each figure are the mean of the estimates among 100 simulations. Lower panels in each figure are Mean Absolute Error (MAE) of the estimates.

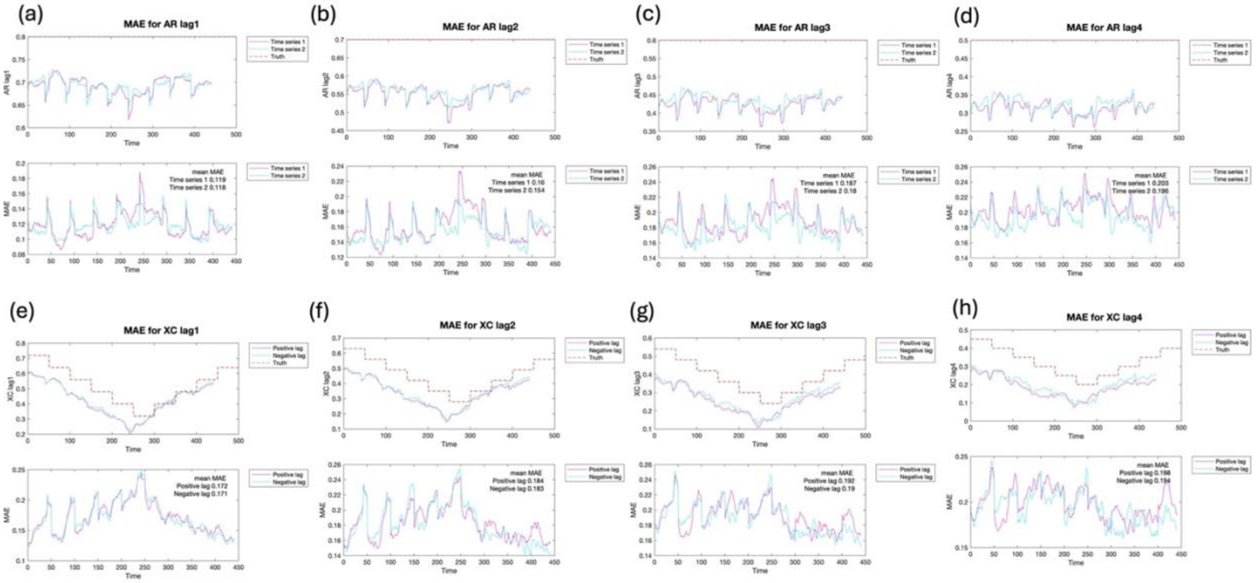


**Supplement Figure 13.** Simulation results of the autocorrelation function (ACF) and cross-correlation function (XCF) estimation using the Fast Fourier Transform (FFT) method. This simulation scenario is Pattern 5 (decreasing followed by increasing) for dynamic correlation with AR7 structure. (a)-(d) ACF at lag 1-4. (e-h) XCF at lag 1-4. Upper panels in each figure are the mean of the estimates among 100 simulations. Lower panels in each figure are Mean Absolute Error (MAE) of the estimates.

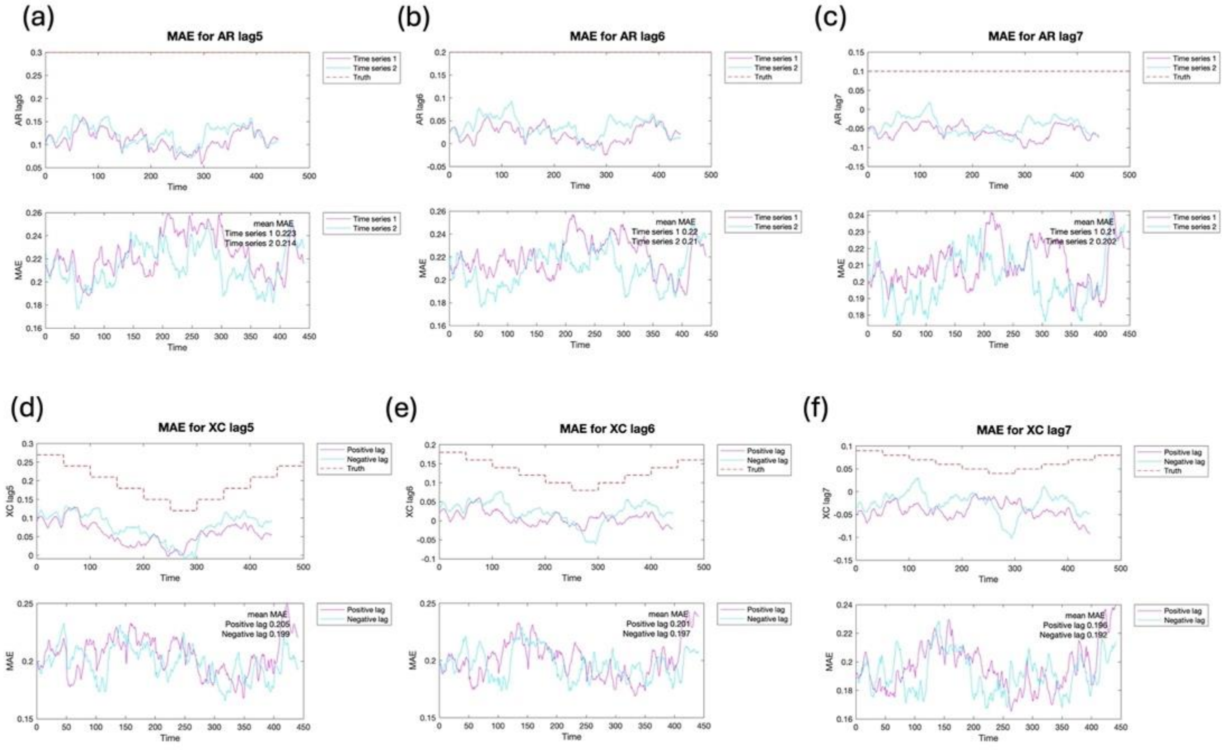


**Supplement Figure 14.** Simulation results of the autocorrelation function (ACF) and cross-correlation function (XCF) estimation using the Fast Fourier Transform (FFT) method. This simulation scenario is Pattern 5 (decreasing followed by increasing) for dynamic correlation with AR7 structure. (a)-(d) ACF at lag 5-7. (e-h) XCF at lag 5-7. Upper panels in each figure are the mean of the estimates among 100 simulations. Lower panels in each figure are Mean Absolute Error (MAE) of the estimates.

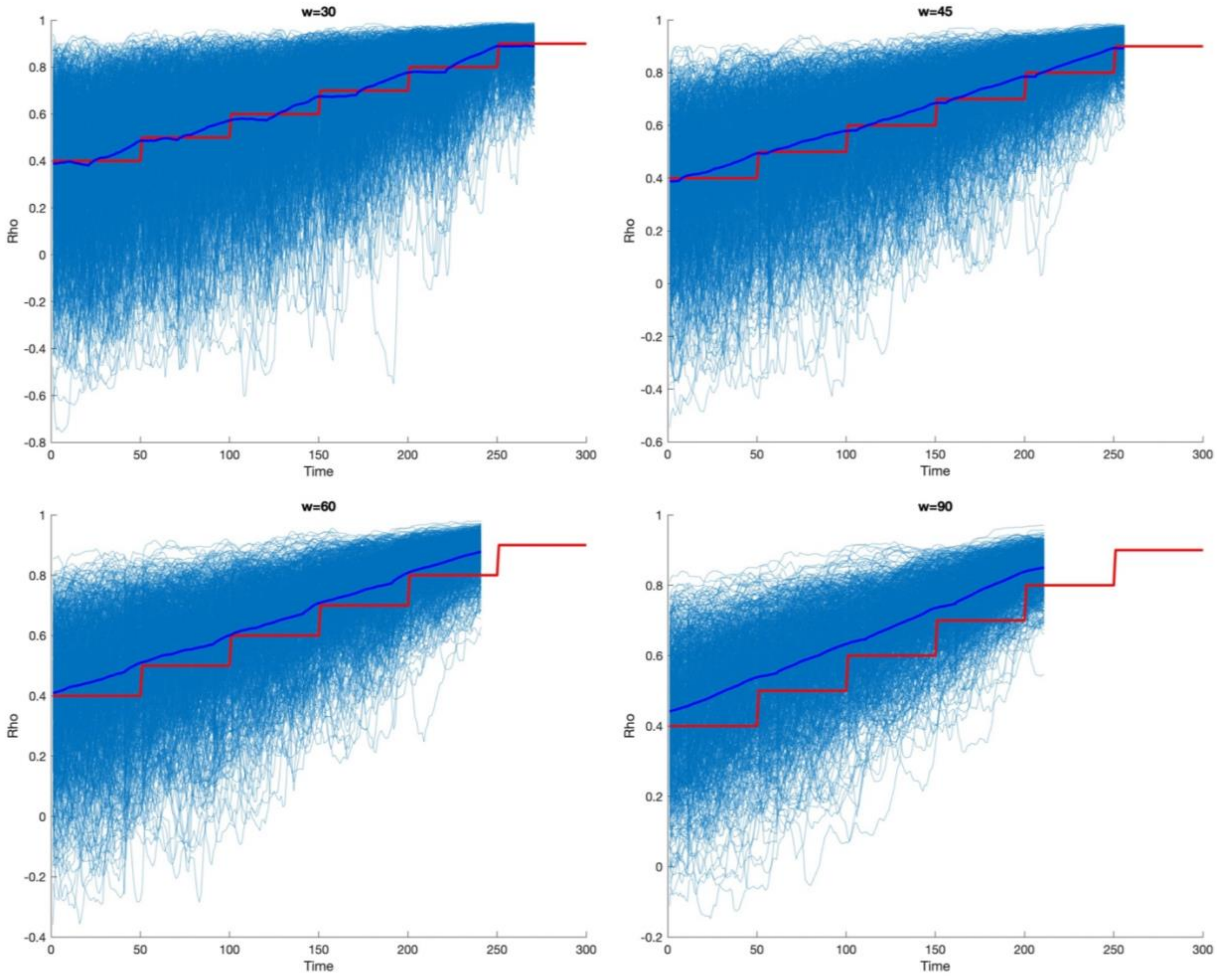


**Supplement Figure 15.** Bias and variance trade-off in sliding window estimation of correlation for different window sizes. The light blue lines represent the estimates from 1000 simulation repetitions. The dark blue line is the mean of the estimates from 1000 simulation repetitions. The red line is the simulation truth.

**Supplement Table 1.** Window size selection for different scenarios with a small tolerance percentage as 5%

| AR | Pattern | Window size for XCF at positive lag 1 ($w_2$) | Relationship to window size for correlation ($w_1$) (Times) |
|---|---|---|---|
| AR4 | 1: Two stages | 40 | 1.33 |
| AR4 | 2: Mono increase | 100 | 1.67 |
| AR4 | 3: Mono decrease | 70 | 1.17 |
| AR4 | 4: Increase and decrease | 100 | 1.67 |
| AR4 | 5: Decrease and increase | 70 | 1.17 |
| AR7 | 1: Two stages | 50 | 1.67 |
| AR7 | 2: Mono increase | 110 | 1.83 |
| AR7 | 3: Mono decrease | 70 | 1.00 |
| AR7 | 4: Increase and decrease | 120 | 1.71 |
| AR7 | 5: Decrease and increase | 80 | 1.33 |

XCF: Cross-correlation function. AR4: Autocorrelation structure with order of 4. AR7: Autocorrelation structure with order of 7.

**Supplement Table 2.** Window size selection for different scenarios with a large tolerance percentage as 15%

| AR | Pattern | Window size for XCF at positive lag 1 ($w_2$) | Relationship to window size for correlation ($w_1$) (Times) |
|---|---|---|---|
| AR4 | 1: Two stages | 30 | 1.50 |
| AR4 | 2: Mono increase | 80 | 1.60 |
| AR4 | 3: Mono decrease | 50 | 1.00 |
| AR4 | 4: Increase and decrease | 80 | 1.60 |
| AR4 | 5: Decrease and increase | 60 | 1.50 |
| AR7 | 1: Two stages | 30 | 1.50 |
| AR7 | 2: Mono increase | 90 | 1.80 |
| AR7 | 3: Mono decrease | 60 | 1.20 |
| AR7 | 4: Increase and decrease | 90 | 1.80 |
| AR7 | 5: Decrease and increase | 60 | 1.20 |

XCF: Cross-correlation function. AR4: Autocorrelation structure with order of 4. AR7: Autocorrelation structure with order of 7.
